# TET enzymes augment AID expression via 5hmC modifications at the *Aicda* superenhancer

**DOI:** 10.1101/438531

**Authors:** Chan-Wang J. Lio, Vipul Shukla, Daniela Samaniego-Castruita, Edahi González-Avalos, Abhijit Chakraborty, Xiaojing Yue, David G. Schatz, Ferhat Ay, Anjana Rao

## Abstract

TET enzymes are dioxygenases that promote DNA demethylation by oxidizing the methyl group of 5-methylcytosine (5mC) to 5-hydroxymethylcytosine (5hmC). Here we report a close correspondence between 5hmC-marked regions, chromatin accessibility and enhancer activity in B cells, and a strong enrichment for consensus binding motifs for basic region-leucine zipper (bZIP) transcription factors at TET-responsive genomic regions. Functionally, Tet2 and Tet3 regulate class switch recombination (CSR) in murine B cells by enhancing expression of *Aicda*, encoding the cytidine deaminase AID essential for CSR. TET enzymes deposit 5hmC, demethylate and maintain chromatin accessibility at two TET-responsive elements, *TetE1* and *TetE2*, located within a superenhancer in the Aicda locus. Transcriptional profiling identified BATF as the bZIP transcription factor involved in TET-dependent *Aicda* expression. 5hmC is not deposited at *TetE1* in activated *Batf*-deficient B cells, indicating that BATF recruits TET proteins to the *Aicda* enhancer. Our data emphasize the importance of TET enzymes for bolstering AID expression, and highlight 5hmC as an epigenetic mark that captures enhancer dynamics during cell activation.

## Introduction

TET proteins (<u>T</u>en-<u>E</u>leven-<u>T</u>ranslocation; TET1, TET2, TET3) are Fe(II)- and α-ketoglutarate-dependent dioxygenases that catalyze the step-wise oxidation of 5-methylcytosine (5mC) to 5-hydroxymethylcytosine (5hmC), 5-formylcytosine (5fC) and 5-carboxylcytosine (5caC)(*1, 2*). Together these oxidized methylcytosine (oxi-mC) bases are intermediates in DNA demethylation, and may also function as stable epigenetic marks. 5hmC, the most stable and abundant product of TET enzymatic activity, is highly enriched at the most active enhancers and in the gene bodies of the most highly expressed genes, and its presence at enhancers correlates with chromatin accessibility. TET proteins regulate several fundamental biological processes including lineage commitment, and play important roles in embryonic, neuronal and haematopoietic development (*3*).

TET proteins, particularly TET2 and TET3, have critical roles in B cell differentiation and malignancy (*2*). We and others have previously shown that deletion of the *Tet2* and *Tet3* with *Mb1-Cre* at early stages of mouse B cell development resulted in impaired light chain rearrangement and developmental blockade, and eventually developed an acute precursor-B-cell-derived leukemia with 100% penetrance (*4, 5*). Inducible deletion of *Tet1* and *Tet2* using *Mx1-Cre* promoted the development of acute lymphoblastic leukemia derived from precursor B cells, and global loss of *Tet1* caused B cell lymphomas with an extended latency (*6*). In humans, *TET2* mutations are frequently observed in Diffuse Large B Cell Lymphoma (DLBCL), a malignancy derived from germinal center B cells (*7, 8*), suggesting that TET proteins may regulate mature B cell function. However, due to the pleiotropic functions of TET proteins, studies of TET-mediated gene regulation are best performed in systems where TET genes are deleted acutely rather than during development.

After their development in the bone marrow, mature B cells migrate to peripheral lymphoid tissues where they encounter antigen and follicular T helper cells in germinal centers, and participate in the generation of functional immune responses(*9*). In germinal centers, B cells undergo Class Switch Recombination (CSR) to replace the constant region of immunoglobulin M (IgM) to other isotypes such as IgG_1_ and IgA for distinct effector functions, and also diversify the variable regions of Ig heavy and light chains for antigen recognition in a process known as somatic hypermutation (SHM). CSR and SHM are both orchestrated by the enzyme AID (Activation-induced cytidine deaminase, encoded by *Aicda*) (*10–12*). AID promotes CSR and SHM by generating DNA double-strand breaks at Ig switch regions and point mutations at Ig variable regions, respectively (*13*). Due to its high mutagenic potential (*14, 15*), AID expression is normally restricted to activated B cells, stimulated either through their B cell receptor and CD40, or through pattern recognition receptors such as TLR4 which binds lipopolysaccharide (LPS).

At the transcriptional level, induction of *Aicda* is tightly controlled by at least 6 conserved *cis*-regulatory elements (*16*). A promoter-proximal region (Region I) and an intronic regulatory element (Region II or E4) function as an enhancer in activated B cells and, in the case of Region II/E4, as a silencer in non-B cells and naïve and memory B cells (*17, 18*). Two additional enhancers, located at −8 kb and +13 kb relative to the *Aicda* transcription start site (TSS), are essential for *Aicda* expression in activated B cells (*19*). The +13kb region (termed CNS-X, E5, or Region III) showed a dramatic increase in histone H3 acetylation after TLR4-dependent activation. Both the −8 kb and +13 kb enhancers are necessary for regulating *Aicda* expression, as shown by studies using bacterial artificial chromosome (BAC) transgenic mice (*17, 19*). Moreover, elements at −26 kb (E1) and −21 kb (E2) are necessary for CSR in CH12F3 B cell line (*20*), suggesting that the concerted action of all enhancers are required for *Aicda* expression. The *Aicda* regulatory elements are occupied by multiple transcription factors, including NFκB, PAX5, E2A, and others (*18, 21*). Transcriptional activation of the *Aicda* gene is also coupled to cell proliferation, although the mechanistic link between these events is not fully understood (*22*).

Here we have investigated the role of TET proteins at a kinetic level during B cell activation, integrating our analyses of TET proteins and 5hmC deposition with data from previous studies that have dissected the transcriptional and epigenetic changes occurring in activated B cells (*20, 23*). We used the *Cre^ERT2^* system for acute gene deletion to avoid secondary effects caused by prolonged *TET* deficiency during cell differentiation. We show that the TET proteins Tet2 and Tet3 are important regulators of CSR in activated murine B cells and that they function by controlling the activation-induced upregulation of AID mRNA and protein. We further demonstrate that transcriptionally, TET proteins act downstream of the bZIP transcription factor Batf, which is also induced during B cell activation with more rapid kinetics than *Aicda*, and binds concomitantly with TET proteins to two TET-responsive elements in the *Aicda* locus that we termed *TetE1* and *TetE2*. Together, our study constitutes a comprehensive analysis of the role of TET proteins in class switch recombination in activated B cells, and provides a detailed analysis of how TET proteins influence cell activation and differentiation.

## Results

### Genome-wide kinetics of 5hmC deposition during B cell activation

We used CMS-IP (*24, 25*) to analyze the kinetics of genome-wide 5hmC distribution in murine B cells activated with LPS and IL-4, a well-characterized *in vitro* system for studying gene regulation (**Fig. 1a**). The vast majority of 5hmC-marked regions (~160,000) were shared between pre- and post-activated B cells (**Fig. 1b**, **Fig. S1a**); of the ~9,500 differentially hydroxymethylated regions (DhmRs) in 72h-activated versus naïve B cells, the majority (8,454) showed increased 5hmC (DhmR^72h-up^) whereas a much smaller fraction showed a decrease (DhmR^down^) (**Fig. 1b,c**). DhmRs were typically located more than 10 kb from the closest transcription start site (TSS) (**Fig. S1b**), and their 5hmC levels progressively changed with time after activation (DhmR^72h-up^; **Fig. 1d**; DhmR^72h-down^; **Fig. S1c**).

**Figure 1.**
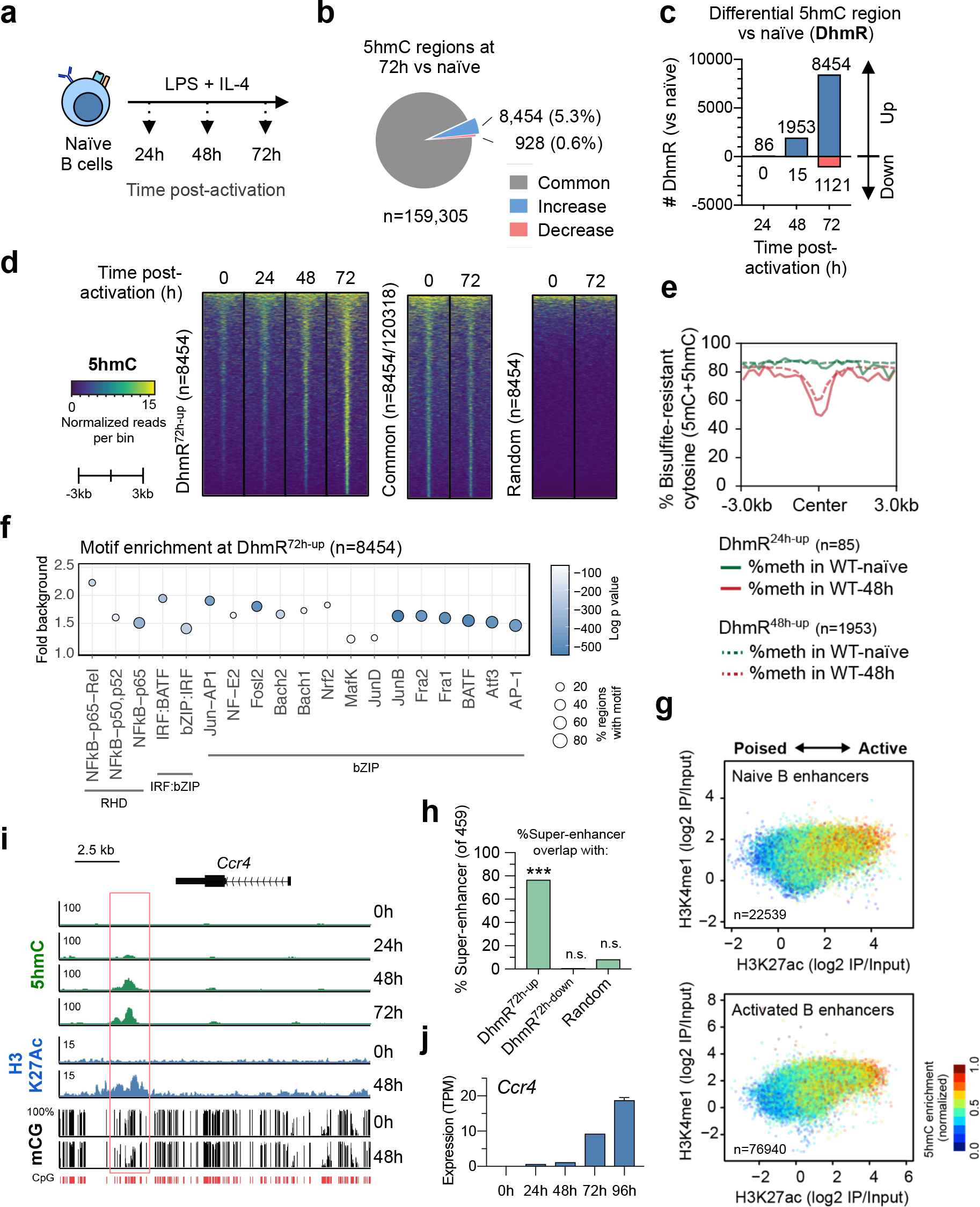
Dynamic changes in 5hmC during B cell activation. **(a)** Flow-chart of experiments. B cells were activated with LPS+IL-4 for the indicated times prior to genome-wide analyses. **(b)** Of a total of 159,305 5hmC-enriched regions in B cells activated for 72h, while most regions (grey, 94.1%) display similar 5hmC levels, 8,454 (blue, 5.3%) show increased 5hmC and 928 (red, 0.6%) show decreased 5hmC relative to naïve B cells. Note that 193 regions represented only in naïve B cells are not shown. **(c)** Number of differential hydroxymethylated region (DhmR) showing increased and decreased 5hmC at respective time points after activation, of a total number of ~160,000 5hmC-marked regions present in naïve and activated B cells (**Fig. S1a**). **(d)** Heatmaps showing the kinetics of 5hmC in the 8,454 regions with increased 5hmC at 72h compared to naïve B cells (*left panels*), but no increase in the same number of 5hmC-marked regions common to naïve and 72h-activated B cells (*middle panels). Right panels*, only a small number of random genomic regions are marked with 5hmC. For a similar analysis of the 1,121 regions that lose 5hmC after 72h of B cell activation, see **Fig. S1c**. 5hmC enrichment is shown as normalized reads per 100 bp bin. **(e)** The 85 and 1,953 regions with increased 5hmC in 24h- and 48h- activated B cells relative to naïve B cells show decreased “methylation” (bisulfite-resistant cytosine, 5mC+5hmC) at their centers 48h after activation. Average methylation was calculated for each 200 bp bin across 6kb. **(f)** Significant enrichment for consensus RHD (NFB), IRF:bZIP, and bZIP transcription factor binding motifs in 8,454 regions DhmR^72h-up^ showing increased 5hmC in 72h-activated relative to naïve B cells. Common 5hmC-enriched regions were used as background for analysis. Y-axis indicates the fold enrichment versus background, circle size indicates the percentage of regions containing the respective motif, and the color indicates the significance (Log_10_ *p* vaule). **(g)** 5hmC is enriched at active (H3K4me1^+^ H3K27Ac^hi^) relative to poised (H3K4me1^+^ H3K27Ac^lo^) enhancers in both activated and naïve B cells. Y and X axes indicate the levels (log_2_) of H3K4me1 and H3K27Ac relative to input, respectively. **(h)** A substantial fraction of super-enhancers (76.7%, 352 of 459) identified by high H3K27Ac enrichment overlap with DhmR^72h-up^ at which 5hmC is increased in activated (72h) relative to naïve B cells. Fisher exact test was used to analyze the significance. ***, *p*<0.01 (*p*=8.9203×10^-266^). n.s., not significant. **(i)** Genome browser view of the *Ccr4* locus (mm10; chr9:114,484,000-114,501,000) as an example of a genomic region marked by increased 5hmC, increased H3K27Ac and decreased CpG methylation in activated compared to naïve B cells. Red track indicates CpGs that were included for analysis based on coverage. **(j)** Kinetics of increase in *Ccr4* mRNA expression (by RNA-seq) in activated B cells. See also Supplementary Fig. 1.

The oxidized methylcytosines produced by TET proteins are known intermediates in DNA demethylation (*2, 26*). To relate changes in 5hmC to changes in DNA methylation, we compared 5hmC distribution in naïve and 72h-activated B cells with published whole-genome bisulfite sequencing (WGBS) data on B cells activated for 48h under similar conditions (*20*). Although WGBS cannot distinguish 5mC and 5hmC (*27*), 5hmC is typically a small fraction (1-10%) of 5mC (*28*), thus we refer to the WGBS signal as “DNA methylation” here. As expected, regions that showed increased 5hmC at 24 and 48h (DhmR^24h-up^, DhmR^48h-up^) also showed decreased DNA methylation at their centers at 48h (**Fig. 1e**). In contrast, only about half (541/1097) of the differentially methylated regions (DMRs) with decreased DNA methylation at 48h (DMR^48h-down^) had increased 5hmC (DhmR^72h-up^, **Fig. S1d-f**), indicating that changes in 5hmC are a more sensitive measure of epigenetic changes in DNA cytosine modification than changes in 5mC.

Motif enrichment analysis of sequences contained in the 8,454 DhmR^72h-up^ peaks (**Fig. 1c**) and the independent set of 1,097 DMR^48h-down^ regions (**Fig. S1d**) showed that both sets of regions were enriched in consensus binding sequences for transcription factors of the NFκB (Rel homology domain, RHD) and basic region-leucine (bZIP) families, as well as for “composite” IRF:bZIP motifs (**Fig. 1f**, **Fig. S1g**) (*29–31*). These motifs were not significantly enriched in the DhmR^72h-down^ regions (<1.5-fold over background; *not shown*), pointing to an association between bZIP and NFκB transcription factors and TET-mediated 5hmC deposition.

To discern the relationship between 5hmC and enhancers, naïve and activated B cell enhancers, defined by H3K4 monomethylation (H3K4me1), were stratified based on the level of H3K27 acetylation (H3K27Ac), a modification that tracks with enhancer activity (*32*). In both sets of enhancers, 5hmC was most highly enriched at active (H3K4me1^+^ H3K27Ac^+^) relative to poised (H3K4me1^+^ H3K27Ac^+^) enhancers (**Fig. 1g**). Moreover, more than 75% of previously identified superenhancers in activated B cells, defined by H3K27Ac, overlapped with at least one DhmR^72h-up^ region (**Fig. 1h**) (*33*). As an example, a 3’ distal element at the *Ccr4* locus showed activation-dependent gain of 5hmC and H3K27Ac, associated with concomitant loss of methylation at specific CpGs and increased mRNA expression (**Fig.s 1i, j**). 5hmC was also associated with accessible chromatin defined by ATAC-seq (*4, 5, 34*) (*see below*), and kinetic analysis of active enhancers, defined as differentially active between naïve and 48h-activated B cells by high accessibility and high H3K27Ac, showed that 5hmC level positively correlated with enhancer activity (**Fig. S1h**). Together our data show that 5hmC modification and DNA demethylation correlates with enhancer activity during B cell activation.

### Comparison of WT and *Tet2/3* DKO B cells identifies TET-responsive regulatory elements

*Tet2* and *Tet3* are the two major TET proteins expressed in B cells (**Fig. 2a**). To evaluate the role of TET proteins in regulating B cell function, we generated mice conditionally deficient in *Tet2* and *Tet3*, using *Cre^ERT2^* and further introduced a *Rosa26-LSL-YFP* cassette to monitor Cre recombinase activity after tamoxifen treatment (LSL: *LoxP*-STOP-*LoxP* cassette in which a strong transcriptional stop is flanked by LoxP sites). *Cre^ERT2^ Tet2^fl/fl^ Tet3^fl/fl^ Rosa26-LSL-YFP* (DKO) and control *Tet2^fl/fl^ Tet3^fl/fl^ Rosa26-LSL-YFP* (WT) mice were treated for 5 days with tamoxifen, after which WT and *Tet2/3* DKO B cells were isolated and activated with LPS and IL-4 (**Fig. 2b**). Both *Tet2* and *Tet3* were efficiently deleted in B cells (**Fig. 2c**), and the YFP^+^ cells showed similar frequencies of mature splenic follicular B cells (**Fig. S2a**).

**Figure 2.**
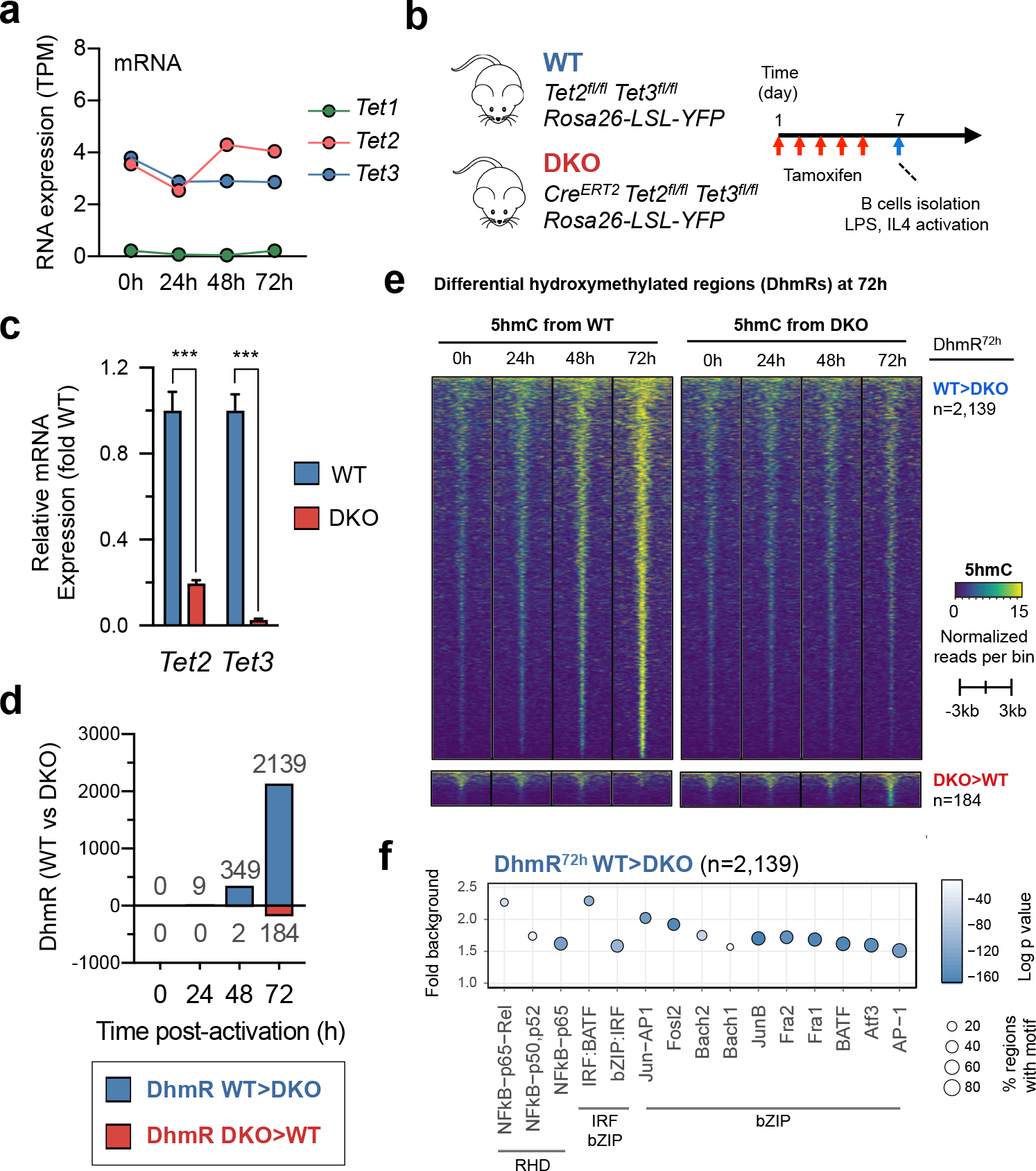
Comparison of 5hmC modification in WT and*Tet2/3* DKO B cells. **(a)** Mean mRNA expression levels for TET family members (from RNA-seq) in WT naïve and activated B cells. TPM, transcript per million. **(b)** Description of mice and flow chart of experiment. **(c)** *Tet2* and *Tet3* are efficiently deleted. *Tet2* and *Tet3* expression in B cells from tamoxifen-treated WT control and *Tet2/3* DKO mice (described in **Fig. 2b**) were analyzed by qRT-PCR. Data were normalized to *Gapdh* within sample and subsequently to the value from WT. Representative of two independent experiments with three technical replicates is shown. *\*\*\*, p*<0.01. **(d)** Number of regions differentially marked with 5hmC (DhmR) between WT and *Tet2/3* DKO B cells as a function of time after activation. **(e)** Heatmaps showing the kinetics of 5hmC enrichment signals from WT (*left panels*) and *Tet2/*3-DKO *(right panels)* at the differentially hydroxymethylated regions (DhmR^72h^) between WT and DKO (72h, **Fig. 2d**). Regions with decreased 5hmC in DKO are shown on top (WT>DKO, n=2,139) and those with increased 5hmC on bottom (DKO>WT, n=184). 5hmC enrichment is shown in normalized reads per 100 bp bin. **(f)** Strong enrichment for consensus RHD (NF_κ_B), IRF:bZIP (IRF:BATF) and bZIP transcription factor binding motifs in the “TET-dependent” regions with decreased 5hmC in 72h-activated *Tet2/3* DKO relative to WT B cells (DhmR^72h-WT>DKO^, n=2,139). Common 5hmC-enriched regions were used as background for analysis. Y-axis indicates the fold enrichment versus background, circle size indicates the percentage of regions containing the respective motif, and the color indicates the significance (Log_10_ *p* vaule). See also Supplementary Fig. 2.

Global 5hmC levels assessed by DNA dot blot were similar in WT and *Tet2/3* DKO B cells prior to activation but showed a perceptible decrease by 48h after activation, as expected since 5hmC is passively lost as a function of cell division (*2, 26*) (**Fig. S2b**). Starting at 48h, 5hmC levels were significantly lower in *Tet2/3* DKO compared to WT B cells, indicating that Tet2 and Tet3 actively oxidize 5mC to 5hmC during B cell activation (**Fig. S2b**). Around 2,300 5hmC-enriched regions were significantly different between WT and DKO at 72h of activation, with substantially more regions gaining 5hmC in control compared to *Tet2/3* DKO B cells at each time point examined (**Fig. 2d, 2e**). Of 2,139 “TET-regulated” DhmRs with higher 5hmC in WT compared to *Tet2/3* DKO B cells (“WT>DKO DhmR”), 2020 (94.4%) significantly overlapped with DhmR^72-up^ (2020/8454, 23.9%). Consistently, “WT>DKO DhmR” were located more than 10 kb from the TSS (**Fig. S2c**), showed decreased DNA methylation at their centers (**Fig. S2d**), and were enriched for NF-κB (Rel homology domain, RHD), bZIP and composite IRF:bZIP motifs (**Fig. 2f**).

### Tet2 and Tet3 regulate immunoglobulin class switch recombination (CSR)

To assess the effect of *Tet2/3* deletion on the antibody response *in vivo*, we treated *Tet2* ^*fl/fl*^ *Tet3*^*fl/fl*^ *Rosa26- LSL-YFP Cre*^*ERT2*^ and control *Tet2* ^*fl/fl*^ *Tet3*^*fl/fl*^ *Rosa26-LSL-YFP* mice for 5 days with tamoxifen, followed by immunization with NP-OVA in the footpads two days later. Germinal center responses in draining popliteal lymph nodes were analyzed by flow cytometry on day 7 post-immunization, gating on YFP^+^ B cells in *Tet2/3* DKO B cells (**Fig. 3a**). The overall percentage of germinal center (GC) B cells (CD19^+^ GL7^+^ Fas^+^) was similar between WT and DKO (**Fig. 3b**, quantified in **Fig. 3c**), indicating that acute deletion of *Tet*2/3 had no immediate impact on the formation of GC B cells. The most striking phenotype, however, was the consistent decrease in CSR from IgM to IgG1 in both total and NP-specific germinal center B cells (**Fig. 3d-f**), demonstrating a role for TET proteins in regulating antibody responses *in vivo*, particularly the CSR.

**Figure 3.**
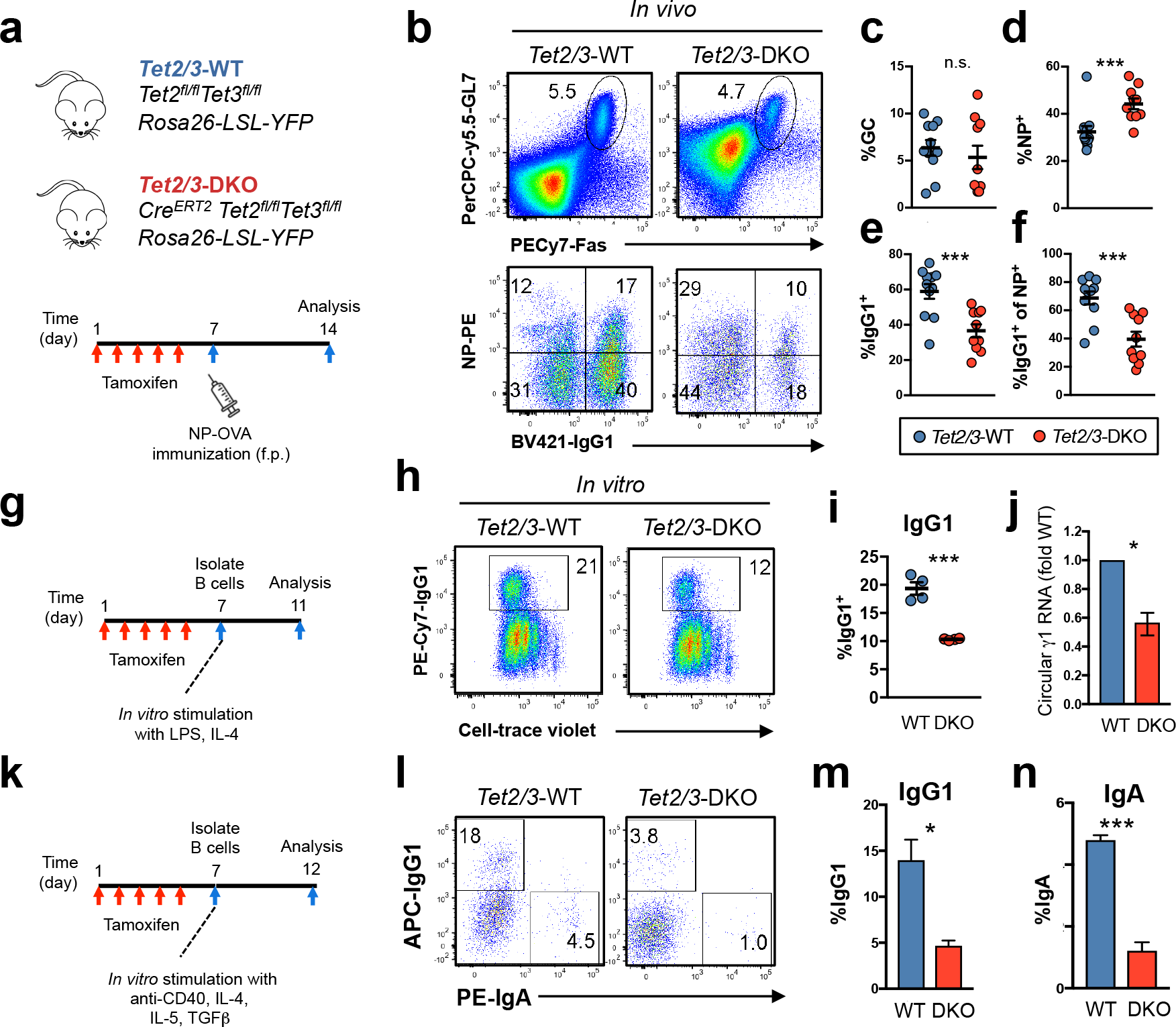
TET proteins facilitate class switch recombination (CSR)*in vitro* and*in vivo*. **(a)** Flow chart of experiment to assess CSR *in vivo*. *f.p.,* foot pad. **(b)** *Upper panels*, flow cytometry plots showing equivalent frequencies of CD19^+^GL7^+^Fas^+^ germinal center B (GCB) cells at the draining popliteal lymph nodes from WT and *Tet2/3* DKO mice after treated with tamoxifen and immunized with NP-OVA as indicated in **(a)**. *Lower panels*, flow cytometry plots showing decreased frequencies of IgG1-switched cells among GCB cells in *Tet2/3* DKO (YFP^+^ GCB-gated) compared to WT mice (GCB-gated). **(c-f)**Quantifications of experiments shown in **(b)**. Data shown are aggregated results from two independent experiments. Means and standard errors are shown. WT, n=11; DKO, n=12. **(g)**Flow chart of experiment to assess CSR (IgG1 switching) *in vitro*. Cells were labeled with Cell-Trace violet and activated for 4 days with LPS (25 ug/mL) and rmIL-4 (10 ng/mL). **(h-i)**Flow cytometry plots (**h**) and quantification of experiments (**i**) show decreased frequencies of IgG1-switched B cells in *Tet2/3* DKO (n=4) compared to WT (n=4) mice. Data were representative from at least three independent experiments. **(j)** Circular γ1 transcript, generated after successful IgG1 switching, was quantified by qRT-PCR and normalized to *Gapdh* and then to the level of WT. Representative of two independent experiment is shown with three technical replicates. **(k)** Flow chart of experiment to assess CSR (IgG1- and IgA-switching) in cell culture. Cells were activated for 5 days with anti-CD40 (1 ug/mL), rmIL-4 (10 ng/mL), rmIL-5 (10 ng/mL), and rhTGFβ (1 ng/mL). **(l-m)** Flow cytometry plots **(l)** and quantification of experiments **(m, n)** showing decreased frequencies of IgG1- **(m)** and IgA-switched cells **(n)** in *Tet2/3* DKO compared to WT cells. Data shown are representative from three independent experiments with three technical replicates. Statistical significance was calculated using unpaired two-tailed *t*-test. n.s., not significant. ***, *p<0.01*. *, *p<0.05*. See also Supplementary Fig. 3.

To determine if the CSR phenotype was B-cell-intrinsic, B cells from tamoxifen-treated mice were labeled with proliferation dye (Cell-trace Violet) and activated with LPS and IL-4 for 4 days (**Fig. 3g**). Consistent with the CSR defect *in vivo*, we noticed a consistent decrease in IgG1 switching in *Tet2/3* DKO B cells activated *in vitro* relative to WT B cells (**Fig. 3h, 3i**). The impaired CSR in *Tet2/3* DKO is not due to Cre activity, as similar result was observed when *Cre^ERT2^ Tet2^+/+^ Tet3^+/+^ Rosa26-LSL-YFP* was used as control (**Fig. S3a**). The defect in CSR was cell-intrinsic, since it was also apparent when congenically-marked WT (CD45.1) and *Tet2/3* DKO (CD45.2) B cells were mixed and co-cultured (**Fig. S3b**). The difference was not due to altered proliferation, which was comparable between WT and *Tet2/3* DKO B cells (**Fig. 3h, Fig. S3c**). Correlating with the decrease in CSR from IgM to IgG1, the expression of circular γ1 transcript was decreased in *Tet2/3* DKO B cells (**Fig. 3j**). Further, CSR to IgA was also decreased in *Tet2/3* DKO relative to WT B cells activated with anti-CD40, IL- 4, IL-5, and TGFβ (**Fig. 3k-3n**). Notably, reconstitution of *Tet2/3* DKO B cells with the enzymatically active catalytic domain of TET2 (Tet2CD) restored CSR from IgM to IgG1 almost to control levels, whereas an enzymatically inactive mutant of Tet2CD (Tet2CD^HxD^) was ineffective (**Fig. S3d-3f**). These results indicate that *Tet2* and *Tet3* are required for optimal CSR both *in vitro* and *in vivo*, and that their catalytic activity is required.

Because the CSR defect in *Tet2/3* DKO B cells was ~50% of control, we asked whether deletion of all three TET proteins might have a more striking effect. We treated *Cre^ERT2^Tet1^fl/fl^ Tet2^fl/fl^ Tet3^fl/fl^ Rosa26-LSL-YFP* and control *Tet1^fl/fl^ Tet2^fl/fl^ Tet3^fl/fl^ Rosa26-LSL-YFP* mice for 5 days with tamoxifen, followed by NP-OVA immunization as in **Fig. 3a**. Consistent with the very low expression of *Tet1* in mature B cells (**Fig. 2a**), the CSR defect in *Tet1/2/3 TKO* was comparable to that observed in *Tet2/3* DKO mice, with decreased IgG1- switched cells (**Fig. S3g-3h**). These results indicate that Tet2 and Tet3 are the major TET proteins that regulate CSR in B cells.

### Tet2 and Tet3 regulate expression of the cytidine deaminase AID

CSR is a highly regulated process and involves multiple pathways, including cytokine signaling and DNA repair (*35*). RNA-seq analysis identified a relatively small number of genes differentially expressed between WT and *Tet2/3* DKO B cells under resting conditions and at different time points after activation (**Fig. S4a, 4b**); among these was *Aicda*, which encodes AID, the activation-induced cytidine deaminase essential for CSR. qRT-PCR analysis confirmed an ~50% decrease in *Aicda* mRNA expression in *Tet2/3* DKO relative to WT B cells at each time point from 48 to 96 hours post-activation (**Fig. 4a, Fig. S4c**), a phenotype reminiscent to the dampened CSR in the case of AID haploinsufficiency (*36, 37*) (**Fig. S4d-i**). The decrease in protein expression was even more profound, as shown by immunoblotting (**Fig. 4b**).

**Figure 4.**
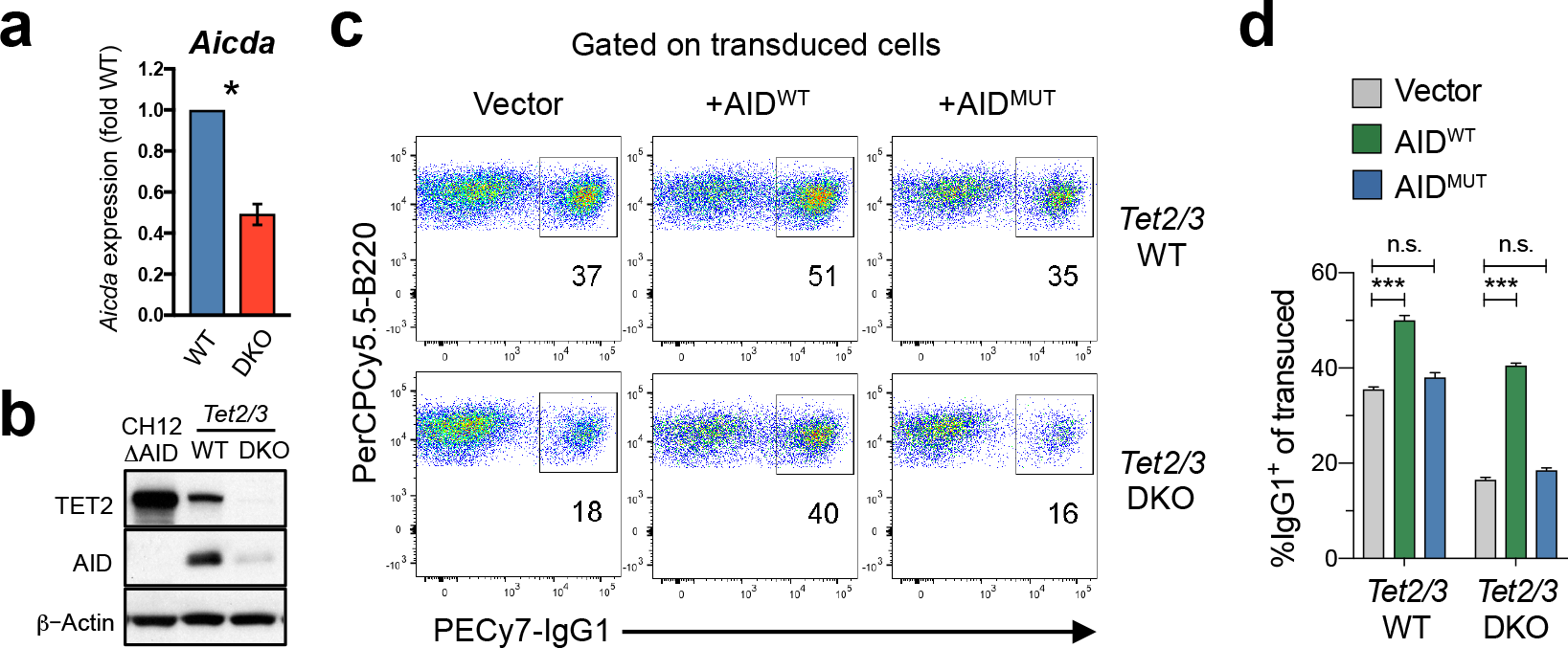
Tet2/3 facilitate CSR by regulating expression of the cytidine deaminase AID. **(a-b) Decreased expression of AID mRNA and proteins in*Tet2/3-***DKO. **(a)**qRT-PCR analysis *Aicda* mRNA expression in WT and *Tet2/3-*DKO B cells activated 4 days with LPS and IL4. *Aicda* expression was normalized to *Gapdh* and then to the level in WT. Result shows ~50% decrease of *Aicda* mRNA expression in *Tet2/3* DKO relative to WT B cells. Data shown are representative of two independent experiments with three technical replicates. *, *p<0.05*. For results of genome-wide RNA sequencing, see **Supplementary Fig. 4**. **(b)** Immunoblotting of whole cell lysates showed a substantial decrease of AID and Tet2 protein expression in *Tet2/3-*DKO relative to WT B cells activated for 4 days. Left lane contains lysate from the AID-KO CH12 B cells as a control for the specificity of anti-AID antibody. β-Actin was used as loading control. Data shown are representative of two independent experiments. **(c-d)** Expression of catalytically active AID restores CSR in *Tet2/3-*DKO. WT and *Tet2/3-*DKO B cells were retrovirally transduced with empty vector expressing Thy1.1 (*left panels*), wild-type AID (AID^WT^, *middle panels*), or catalytically inactive AID (AID^MUT^, *right panels*). Cells were gated on live transduced B cells (CD19^+^ Thy1.1^+^). Representative flow cytometry plots **(c)** and quantification **(d)** are shown. Data shown are representative of three independent experiments. n.s., not significant. ***, *p<0.01*. See also Supplementary Fig. 4.

To determine if the decrease in AID expression was fully responsible for the CSR defect, we expressed WT and catalytically inactive AID in WT and *Tet2/3* DKO B cells via retroviral transduction. Prior to transduction, B cells from WT and DKO mice were stimulated for 24 hours with LPS and F(ab’)_2_ anti-IgM, a stimulation condition in which CSR was inhibited and AID expression was delayed (*38*). CSR was induced with LPS and IL-4 24h post-transduction. We argued that if Tet2 and Tet3 regulated additional major process(es) downstream of AID, re-expressing AID in *Tet2/*3 DKO would only have a marginal effect on the observed CSR phenotype. In fact, retroviral expression of catalytically active AID (AID^WT^) largely rescued the CSR defect in *Tet2/3* DKO B cells; catalytic activity was required as expression of a catalytically inactive AID mutant (AID^MUT^) had no effect on CSR (*bottom panels*; **Fig. 4c, 4d**). Similar to previous observations (*12*), expression of AID in WT cells also increased the frequency of IgG1^+^ cells (*top left and middle panels*; **Fig. 4c, 4d**). Despite their importance in *Aicda* expression, Tet2/3 were not required for the expression of μ and γ1 germline transcripts that are essential for CSR (**Fig. S4j**). These data suggest that the bulk of the CSR defect in *Tet2/3* DKO B cells can be attributed to the decrease in expression of *Aicda* mRNA and AID protein, leading us to test the hypothesis that TET proteins control *Aicda* expression through distal regulatory element(s) of the *Aicda* gene.

### Identification of TET-responsive regulatory elements in the *Aicda* locus

Multiple conserved regulatory elements influence *Aicda* expression (**Fig. S5a**), including an intronic region located 26 kb 5’ of the *Aicda* TSS, within the adjacent *Mfap5* gene; two intergenic regions between the *Mfap5* and *Aicda* genes (located at −21kb and −8 kb 5’ of the *Aicda* TSS respectively); a region in the first intron of *Aicda*; and a region located 3’ of the *Aicda* gene. Deletion of any of the above elements, either in the endogenous locus or in the context of BAC transgenes, dramatically decreased the expression of *Aicda* in activated B cells (*17–20*), indicating that they have essential, non-redundant regulatory (enhancer) function and that their concerted action is necessary for *Aicda* induction. The *Aicda* 5’ enhancer at −26 kb in the *Mfap5* gene, the intergenic 5’ enhancers, and the intron 1 enhancer noticeably gain H3K27Ac and lose 5mC upon activation, and have been collectively termed the *Aicda “*superenhancer” (*20, 33*) (**Fig. 5a**, *middle and bottom tracks*).

**Figure 5.**
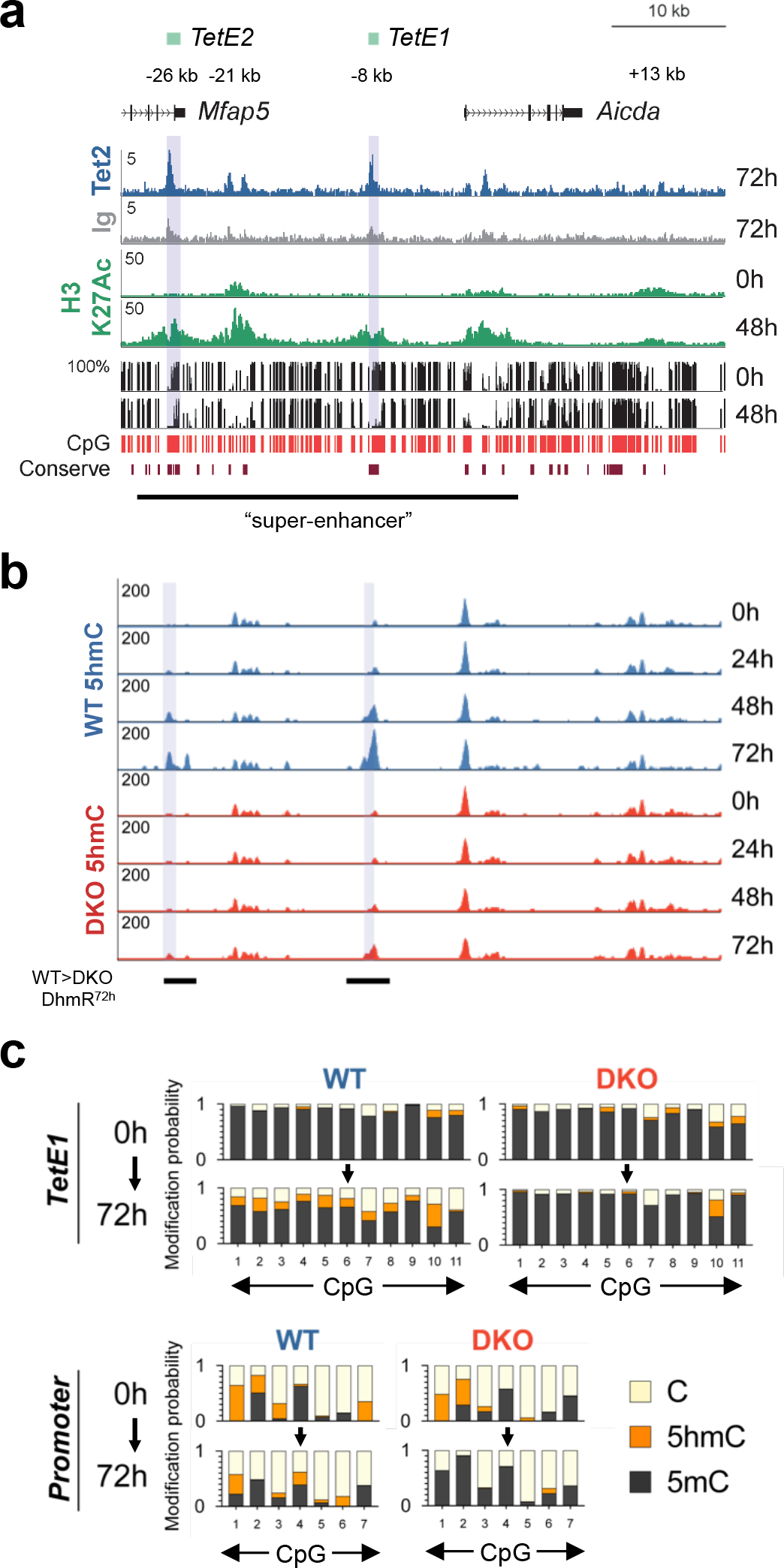
Tet2 and Tet3 control*Aicda* expression via TET-responsive elements *TetE1* and *TetE2*. Diagram shows two conserved TET-responsive elements *TetE1* and *TetE2* at the 5’ of the *Aicda* gene (labeled with green rectangles and grey shades). **(a)** *Top two tracks*. ChIP-seq analysis showed that Tet2 (*blue track*) specifically bound to multiple elements in the *Aicda* locus (mm10; chr6:122,523,500-122,576,500) after activation when compared to Ig control (*grey track*). *Middle tracks* (green). Increased H3K27 acetylation at the upstream and intronic regulatory elements of *Aicda* after activation. *Bottom tracks*. Activation induced DNA demethylation at *TetE1* and *TetE2*. Whole genome bisulfite sequencing (WGBS) showing DNA methylation (5mC+5hmC) in naïve and 48h-activated B cells (*mCG, black tracks*). CpGs included in the analysis are indicated by red lines (*red track*). Bottom track indicates the conserved DNA elements among placental animals (“Conserve”). Previously identified super-enhancer is indicated. For Tet2 and Ig, scales indicate per 10 millions reads; for H3K27Ac, quantile-normalized reads; for bisulfite sequencing, percentage of bisulfite-resistant cytosine. **(b) Activation induced Tet2/3-dependent 5hmC deposition at *Aicda* distal elements.**. WT and *Tet2/3-*DKO B cells were activated as in **Fig. 3g** with LPS and IL-4 as a function of time. DNA was purified and 5hmC enrichment was detected by CMS-IP (see *Materials and Methods*). Significant differential 5hmC-enriched regions between WT and DKO after 72h-activation were indicated at the bottom (WT>DKO DhmR^72h^). Scales indicate quantile-normalized reads. **(c) Tet2/3 deposit 5hmC and demethylate*Aicda* TET-responsive element*TetE1* and promoter.** CpG modifications (5hmC, 5mC, and C) at *TetE1* (*top panels*) and *promoter* (*bottom panels*) were analyzed by oxidative bisulfite sequencing (oxBS-seq; *Materials and Methods*) using DNA isolated from WT and *Tet2/3-*DKO B cells before and after activation. Although 5hmC and 5mC can be distinguished by oxBS-seq, unmodified C and minuscule amount of fC and caC were recognized as “C”, all of which are sensitive to deamination by bisulfite treatment. See also Supplementary Fig. 5.

To determine if these regulatory elements were detectably occupied by Tet2 upon activation, we performed chromatin immunoprecipitation-sequencing (ChIP-seq) for Tet2 in B cells at 72h following activation. Indeed, each of these elements was occupied by Tet2 in 72h-activated B cells (**Fig. 5a**, *top two tracks*). Among these, the *Mfap5* intronic region and the intergenic region, located at −26 kb and −8 kb 5’ of the TSS were clearly “TET-regulated”: their 5hmC levels increased after activation of WT B cells, and this increase was significantly diminished in *Tet2/3* DKO B cells (**Fig. 5b**), placing them in the category of WT>DKO DhmRs (**Fig. 2d, 2e**). We have termed these enhancers *TetE1* (−8 kb) and *TetE2* (−26 kb) respectively; *TetE1* is potentially the prime target for Tet2/3 due to its larger gain of 5hmC after activation (**Fig. 5b**).

To confirm the importance of *TetE1* in *Aicda* regulation, we deleted the enhancer using CRISPR in CH12F3 cells, a B cell line that can class-switch from IgM to IgA upon activation with anti-CD40/IL-4/TGFβ (**Fig. S5b, 5c**). We tested four clones with homozygous deletions; all showed decreased expression of *Aicda* mRNA, and in three of these, there was almost no detectable CSR (**Fig. S5d, 5e**), confirming a previous report in the context of a BAC transgene that *TetE1* was essential for *Aicda* expression (*17*).

B cell activation induces strong DNA demethylation (loss of WGBS signal) at *TetE1* (**Fig. 5a**). Since bisulfite sequencing does not distinguish 5mC and 5hmC, we used oxidative bisulfite sequencing (oxBS-seq) to assess the levels of 5mC, 5hmC and unmodified C at *TetE1* in WT and *Tet2/3-*DKO cells (neither BS-seq nor oxBS-seq distinguish unmodified C from 5fC and 5caC, but these modified bases are ~10-fold and ~100-fold less abundant than 5hmC (*2*)). Consistent with the fact that global 5mC and 5hmC levels are similar between WT and *Tet2/3* DKO B cells prior to activation (**Fig. S2b**), CpGs in both *TetE1* and the *Aicda* promoter displayed similar levels of 5mC and 5hmC prior to activation (**Fig. 5c**, *compare 0 h panels*). At 72h following activation, however, there was a substantial increase in both 5hmC and unmodified C in WT B cells, whereas *Tet2/3* DKO B cells displayed little or no 5hmC and remained significantly methylated (**Fig. 5c**, *compare 72h panels*). These results indicate that Tet2 and Tet3 regulate *Aicda* expression by binding to and depositing 5hmC at both 5’ enhancers, *TetE1* and *TetE2*.

### Tet2 and Tet3 maintain chromatin accessibility at the *Aicda* Tet-responsive elements *TetE1* and *TetE2*

Transcriptional regulatory regions are typically found in accessible regions of chromatin (*39*), and were shown to be marked by 5hmC during T and B cell development (*4, 34*). To assess the dynamics of changes in chromatin accessibility, we performed ATAC-seq in B cells stimulated with LPS and IL-4. Activated B cells displayed progressive chromatin remodeling, with thousands of regions showing changes in accessibility compared to naïve B cells, ranging from 15,529 differentially accessible regions (DARs) at 24h to 28,106 DARs at 72h (**Fig. 6a**). Regions with increased 5hmC after activation (DhmR^72h-up^) also showed increased chromatin accessibility following activation, and vice versa (**Fig. S6a**, *blue box-and-whisker plots* and *not shown*).

**Figure 6.**
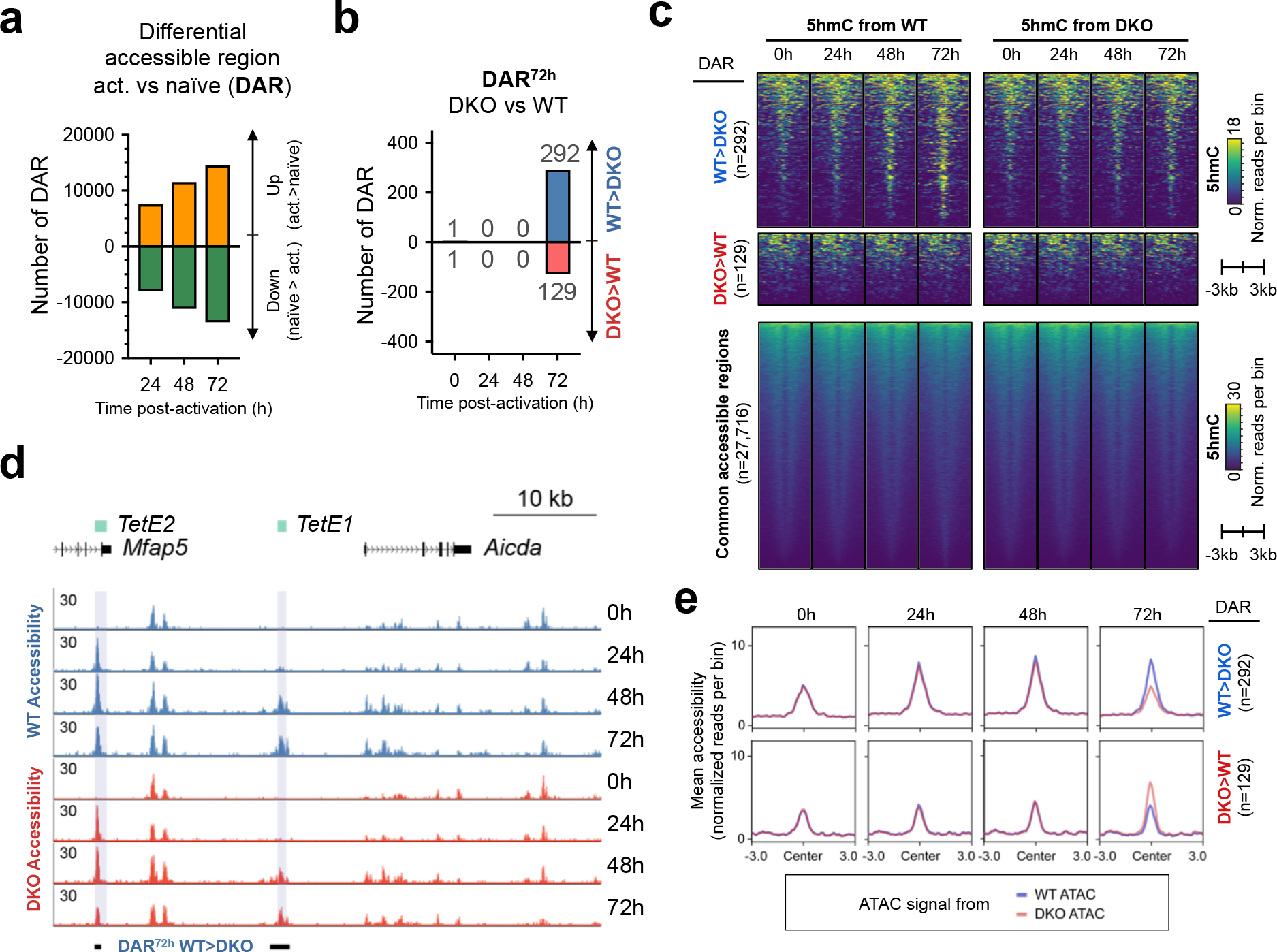
Tet proteins sustain enhancer accessibility. **(a) B cell activation induced global accessibility changes**. WT B cells were activated with LPS and IL-4 as a function of time and the chromatin accessibility was profiled by ATAC-seq. Numbers are shown for the differential accessible regions (DARs) between activated (act.) and naïve B cells with FDR < 0.05 and fold change above log_2_(1.5) or below log_2_(0.67). **(b) Loss of TET proteins decreased accessibility at later time point**. Numbers of DARs between WT and *Tet2/3-*DKO B cells activated across time points are shown. Note that the difference between WT and DKO was minimal at time points earlier than 72h. DARs were selected based on FDR < 0.05 and fold change above log_2_(1.5) or below log_2_(0.67) **(c) Tet2/3-dependent accessible regions are hydroxymethylated**. Heatmaps show the kinetics of 5hmC modification at the differential accessible regions (DARs) between WT and *Tet2/3-*DKO. Regions that are more accessible in WT (WT>DKO, n=292), less accessible in WT (DKO>WT, n=129), and commonly accessible (n=27,716) are shown on top, middle, and bottom panels, respectively. Note that the WT>DKO DARs show progressive 5hmC enrichment only in WT (*top left panels*) but not in DKO (*top right panels*), demonstrating the 5hmC modification at these regions is *Tet2/3*-dependent. The DKO>WT DARs (n=129) and common regions (n=27716) show no apparent difference between naïve and activated B cells and between 5hmC from WT and DKO B cells. Note that the length of the commonly accessible regions is not to scale compared to DARs. 5hmC enrichment is shown as normalized reads per 100 bp bin. **(d) Tet2 and Tet3 maintain chromatin accessibility at*Aicda* Tet-responsive elements*TetE1* and*TetE2***. Genome browser view of ATAC-seq data showing the accessibility profile at *Aicda* locus in WT (*blue, top 4 tracks*) and DKO B cells (*red, bottom 4 tracks*). Note that *TetE1* and *TetE2* were among the DAR at 72h (DAR^72h^ WT>DKO) as indicated at the bottom. Coordinate for locus is chr6:122,523,500-122,576,500 (mm10). **(e)** Plot of mean chromatin accessibility at the DARs between WT and *Tet2/3-*DKO B cells after 72h-activation (as in **(c)** top and middle panels). *Top panels*, WT>DKO DARs (n=292); *bottom panels*, DKO>WT DARs (n=129). Y-axes indicate the mean ATAC signals (normalized ATAC-seq reads per 100bp bin) from WT (blue line) and DKO (red line) B cells activated as indicated. Note that the difference between WT and DKO is apparent at 72h. See also Supplementary Fig. 6.

To understand the relationship between TET function and chromatin accessibility, we performed ATAC-seq on WT and *Tet2/3* DKO B cells activated as in **Fig. 3g**. Of a total of ~28,000 accessible regions (**Fig. 6b, 6c**), only a minor fraction (~1.5%; 421 of 28137) showed significant changes in accessibility between WT and *Tet2/3* DKO B cells and the differences were observed late, at 72h following activation (**Fig. 6b-d**). Of the 292 potentially TET-regulated DARs, defined as showing decreased accessibility in *Tet2/3* DKO compared to WT B cells (WT>DKO DARs), the majority were located distal to the TSS (**Fig. S6b**) and showed a Tet2/3-dependent increase in 5hmC (**Fig. 6c**, **Fig. S6c**, *upper panels*). In contrast, the 129 DKO>WT DARs that were less accessible in WT compared to *Tet2/3* DKO B cells, and the 27716 commonly accessible DARs, were present in both TSS-proximal and -distal regions, and did not show significant changes in 5hmC (**Fig. 6c**, *middle and lower panels;* **Fig. S6b, 6c**). Analysis of DNA methylation at 48h post-activation showed that WT>DKO DARs were further demethylated after activation, whereas DKO>WT DARs were already substantially demethylated in naïve B cells and showed no further changes after activation (**Fig. S6d**). Moreover, WT>DKO DARs resembled DhmR^72h-up^ (**Fig. 1f**) and WT>DKO DhmRs (**Fig. 2f**) in their enrichment for bZIP and BATF:IRF motifs (**Fig. S6e**).

Focusing on the *Aicda* locus, we found that activation was associated with increased accessibility at the *Aicda* enhancers *TetE1* and *TetE2* (**Fig. 6d**). The 5hmC modification continuously increased at these two elements until 72h, with a higher level of deposition at *TetE1* (see **Fig. 5b**). In contrast, the time course of increase in chromatin accessibility was quite different at the two enhancers (**Fig. 6d**): *TetE2* showed a rapid increase in accessibility apparent in both WT and *Tet2/3* DKO B cells at 24 h following activation, whereas the time course of increase in *TetE1* accessibility was slower, matching that of 5hmC deposition (compare **Fig.s 5b** and **6d**). Consistent with the increased accessibility, several chromatin remodelers and histone acetyl-transferases including Brg1, Chd4, p300, and to a lesser extent, Gcn5, were recruited to *TetE1* and *TetE2* in 24h activated B cells (**Fig. S6f**). Interestingly, we noticed a slight decrease in chromatin accessibility at *TetE1* and *TetE2* in *Tet2/3* DKO B cells compared with WT cells at 72 hours, suggesting that TET proteins are important for maintaining the accessibility at these enhancers (**Fig. 6e**). Together these data point to a consistent link between bZIP-family transcription factors, TET catalytic activity and chromatin accessibility that is explored in greater detail below

### Batf acts upstream of TET at*Aicda* enhancers

Before enhancers are established during development, cell lineage specification or activation, certain key transcription factors bind to nucleosome-associated regions and recruit chromatin remodelling complexes and histone modifying enzymes to create active enhancers (*32*). Chromatin remodelling complexes establish accessible (nucleosome-depleted) regions, while the histone acetyltransferase p300/CBP deposits the “active” histone modification H3K27Ac (*32*). To identify potential pioneer transcription factors for the *Aicda* locus, we took advantage of our previous motif enrichment analyses (**Fig.s 1f**, **2f**; **Fig. S1g**). We had observed strong enrichment for consensus binding motifs for bZIP transcription factors, at regions that progressively gained 5hmC as a function of activation (DhmR^up^, **Fig. 1f**), regions that lost DNA methylation upon activation (DMR^48h-down^; **Fig. S1g**), regions with higher 5hmC in WT compared to *Tet2/3* DKO B cells (WT>DKO DhmRs, **Fig. 2f**), and regions with higher accessibility in WT versus *Tet2/3* DKO B cells (DAR^72h^ WT>DKO; **Fig. S6e**).

Based on these data, we focused on bZIP transcription factors expressed in activated B cells. Consistent with previous observations (*40, 41*), *Batf* was expressed at low levels prior to activation but was highly expressed from 24h to 96h following activation (**Fig. 7a**). This increase in *Batf* expression preceded that of *Aidca*, as expected if Batf regulated *Aidca* mRNA induction (**Fig. S7a**). In contrast, expression of *Bach1* and AP-1 (Fos and Jun) family members was either low throughout (*Fosl1, Fosl2, JunD, Bach1*), or moderate to high in unstimulated B cells and strongly reduced by 24h (*Fos, FosB, Jun, JunB*) or 72h (*Bach2*) of activation (**Fig. S7b-d**). Given the kinetics, we examined the importance of Batf in subsequent experiments.

**Figure 7.**
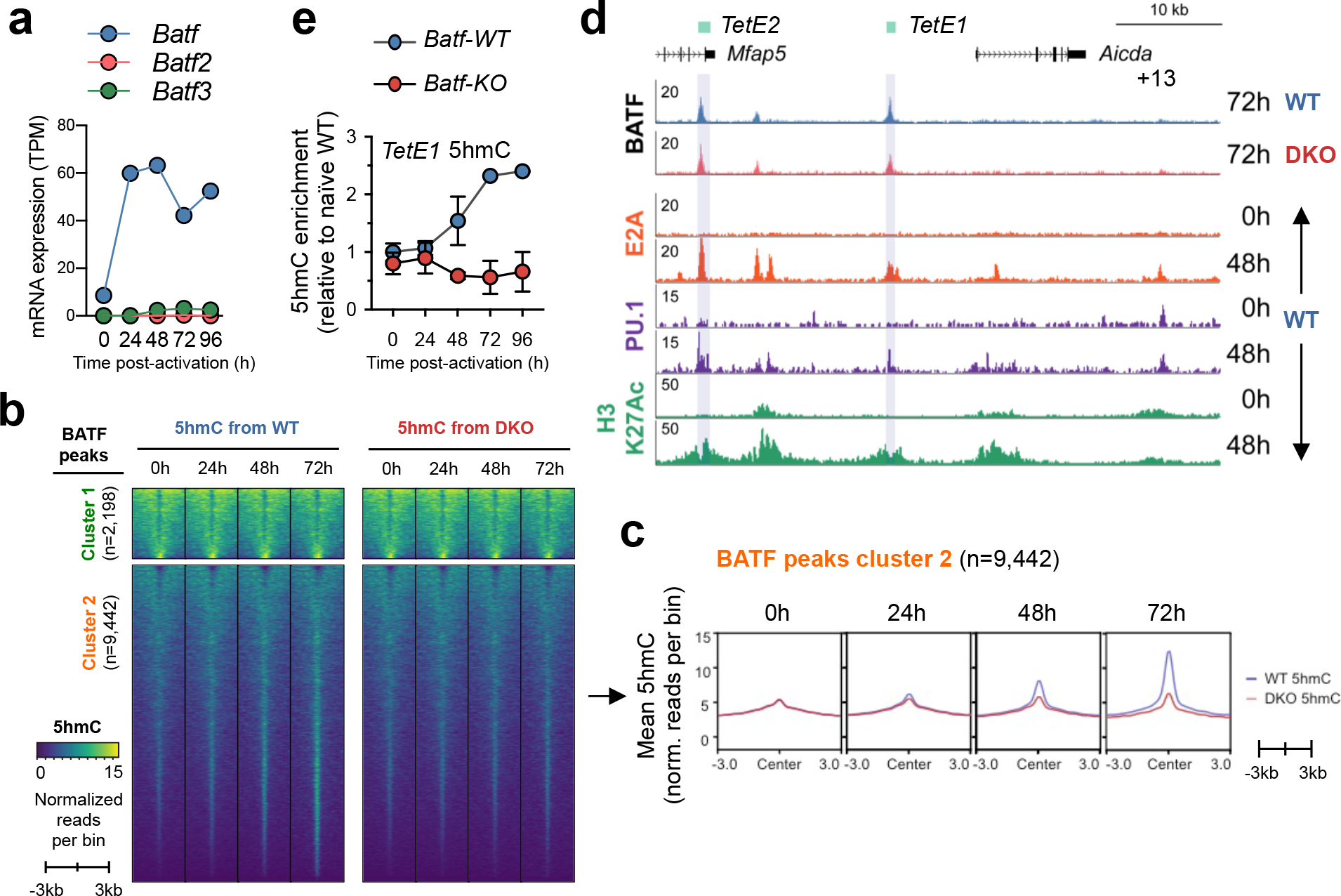
BATF recruits Tet proteins to hydroxymethylated*Aicda* Tet-responsive element*TetE1*. **(a)** Mean mRNA expression level of *Batf* family members (*Batf1-3*) in B cells activated with LPS and IL-4 as a function of time. Data shown are from RNA-seq with two independent replicates. TPM, transcript per million. **(b-c) BATF binding correlates with 5hmC-enrichment**. WT BATF peaks (n=11,640) were divided into two clusters based on the pattern of 5hmC distribution. **(b)** Cluster 1 (n=2,198; *top panels*) showed a broad 5hmC distribution, with the 5hmC level remained unchanged after activation and in the absence of Tet2/3 (*upper panels, compare “5hmC from WT” to “5hmC from* DKO*”*). In contrast, a substantial portion of regions in cluster 2 (n=9,422) showed a progressive Tet-dependent 5hmC modification after activation (*lower panels*) and is further illustrated in **(c)** as line plots. Data shown are mean enrichment per 100 bp bin. **(d) Recruitment of BATF and other transcription factors to*Aicda* enhancers**. *Upper two tracks*, genome browser view of BATF-binding in 72h-activated WT (*blue*) and *Tet2/3-*DKO B cells (*red*) at the *Aicda* locus. Note that the major BATF-binding sites are at *TetE1* and *TetE2,* and the loss of *Tet2/3* has no significant effect on BATF recruitment (compare WT and DKO; also see **Supplementary Fig. 7f**; two independent experiments). Activation also induced E2A and PU.1 binding to *Aicda* enhancers (orange and purple tracks). Coordinate for locus is chr6:122,523,500-122,576,500 (mm10). See also Supplementary Fig. 7. **(e) BATF is required for 5hmC modification at*TetE1***. B cells were isolated from WT and *Batf-KO* and activated with LPS and IL-4 for 4 days. 5hmC modification at *TetE1* was quantified using AbaSI-qPCR (*see Materials and Methods*).

Batf is essential for controlling both T and B cell function during humoral responses (*40, 41*). Importantly, Batf-KO B cells are defective in CSR (**Fig. S7e**) (*40, 41*), thus resembling *Tet2/3* DKO B cells (**Fig. 3**). Genome-wide analysis of Batf binding by ChIP-seq in 72h-activated WT and *Tet2/3*-DKO B cells showed very few overall differences (**Fig. S7f**), indicating that Batf functioned upstream or independently of TET enzymes. Nevertheless, one of two distinguishable sets of Batf ChIP-seq peaks (Cluster 2 in **Fig. 7b**; n=9,442) was clearly TET-regulated, since the peaks in this cluster showed a progressive Tet2/3-dependent increase in 5hmC with time after activation (**Fig. 7b, c**). In contrast, the majority of Batf peaks in Cluster 1 (n=2,198) showed no significant activation-dependent increase in 5hmC (**Fig. 7b**; **upper panel**).

Batf bound strongly at the *TetE1* and *TetE2* enhancers in the *Aicda* locus, and to a lesser extent to the −21 kb intergenic enhancer located between *TetE1* and *TetE2* (**Fig. 7d**, top tracks). This binding pattern resembles that of Tet2 (**Fig. 5a**), as well as that of E2A and PU.1 (**Fig. 7d**), three transcriptional regulators already expressed in unstimulated B cells (**Fig. S7g, 7h**; **Fig. 2a**)(*21, 42, 43*). Moreover, BATF as well as JUNB associated with TetE1 and TetE2 in a human B cell lymphoblast (**Fig. S7i**), suggesting the binding of BATF is evolutionarily conserved. To determine whether Batf acted upstream of TET proteins (for instance by recruiting Tet2 and Tet3 to Batf-bound regulatory regions), we analysed 5hmC deposition at the Batf- and TET-responsive element *TetE1* in WT and Batf-deficient B cells. We found unambiguously that the absence of Batf abolished activation-induced hydroxymethylation at *TetE1* (**Fig. 7e**), indicating that *Batf* is required for TET-mediated hydroxymethylation at this *Aicda* enhancer. Our results are consistent with the hypothesis that Batf recruits Tet2 and/or Tet3 to *TetE1* and *TetE2* and facilitates *Aicda* expression by promoting 5hmC modification and DNA demethylation at these upstream *Aicda* enhancers.

## Discussion

TET proteins (TET1, TET2 and TET3) oxidize 5mC to 5hmC, a stable epigenetic mark that is the most abundant of the three oxi-mC intermediates for DNA demethylation. Due to the pleiotropic effects of TET proteins in cells, it has been challenging to address the specific roles of TET proteins in mice with prolonged TET deficiency. Here, to circumvent this issue, we used the inducible tamoxifen-*Cre^ERT2^* system to delete *Tet2* and *Tet3* in mature B cells, a well-established system for the molecular analysis of gene regulation during cell activation. Our data show clearly that Tet2 and Tet3 – the major TET proteins expressed in B cells – are required for efficient class switch recombination (CSR) both *in vivo* and in cultured cells. A primary mechanism involves TET-mediated regulation of the expression of *Aicda*, the essential DNA cytosine deaminase for CSR. The activation-induced transcription factor BATF, potentially with other transcription factors, recruits TET proteins to two major TET-responsive regulatory elements that we have newly defined in the *Aicda* locus, *TetE1* and *TetE2*. Tet2 and Tet3 convert 5mC to 5hmC at these regulatory elements, leading to DNA demethylation, sustaining enhancer accessibility and augmenting *Aicda* expression.

The biological consequences of TET loss-of-function are determined by several factors: the time course of *Tet2* and *Tet3* gene deletion, the stability of Tet2 and Tet3 mRNA and protein, and the rate of cell division which determines the rate of passive (i.e. replication-dependent) dilution of 5hmC. At each cell division, hemi-methylated CpGs are recognised by the maintenance UHRF1/DNMT1 DNA methyltransferase complex and converted back to symmetrically methylated CpGs, whereas hemi-hydroxymethylated CpGs are ignored and so are diluted by half (*1, 2*). Consequently, 5hmC is present at comparable levels in quiescent (non-dividing) WT and *Tet2/3* DKO B cells, thus enabling us to study the effects of acute TET deletion in activated, proliferating B cells. The progressive replication-dependent loss of 5hmC and consequent dilution of both 5mC and 5hmC is likely to be required for optimal gene expression, explaining the long-standing observation that the induction of *Aicda* expression during B cell activation, and the induction of cytokine genes during Th2 differentiation, are both tightly coupled to cell division (*22, 44*).

An optimal level of AID is crucial to maintain the necessary balance between effective antibody immune responses and unintentional C>T mutations caused by AID-mediated DNA cytidine deamination. Indeed, while *Aicda* haploinsufficiency results in dampened antibody responses (*36, 37*) (**Fig. S4e-j**), uncontrolled *AICDA* expression is associated with B cell malignancies (*45*). Thus, the level and activity of AID are meticulously controlled by diverse mechanisms including a tight transcriptional regulatory program (*16*). At the *Aicda* locus, at least six regulatory elements have been identified (**Fig. S5a**); five of them, located at distances ranging from −29 to +5 kb relative to the TSS, are collectively termed the *Aicda* superenhancer (*20, 33*). The enhancers at −26, −21, −8, +13 kb are all necessary for inducing *Aicda* expression in activated B cells, based on deletion of individual enhancers in mice and the CH12 B cell line (*17, 18*). Notably, even in naïve B cells where *Aicda* is not expressed, the *Aicda* promoter is already highly enriched in 5hmC and the −21, intronic, +13 kb *Aicda* enhancers display 5hmC and H3K27Ac (**Fig. S5a**). In fact, the 5hmC modification at the −26, −21, +13 kb *Aicda* enhancers is apparent as early as the pro-B cell stage of B cell development (*4*), suggesting that TET-mediated 5hmC modification acts to “bookmark” regulatory elements necessary for proper gene expression in progeny cells after activation.

The vast majority of 5hmC-marked regions are present in common between naïve and activated mature B cells (this study), and between WT and TET-deficient iNKT cells (*34*), supporting the hypothesis that most 5hmC-marked regions in any given cell type were laid down during prior developmental stages and thus are constitutively modified. In contrast, activation-induced 5hmC modification occurs at only a few distal elements in B cells (**Fig. 1c**), and 5hmC levels at these elements correlate strongly with activation-induced increases in enhancer activity defined by H3K27Ac (**Fig. 1g**) (*46*). Moreover, the majority of previously described B cell superenhancers (*20, 33*) harbor at least one activation-induced, 5hmC-modified regulatory element (**Fig. 1h**). In the particular case of *Aicda*, we identified activation-induced 5hmC modification at two major TET-responsive elements, *TetE1* and *TetE2*, both part of a superenhancer cluster located 5’of the *Aicda* gene **(Fig. 5a)**(*20, 33*). 5hmC modification at these elements was apparent by 48h (**Fig. 5b**), preceding the dramatic upregulation of *Aicda* mRNA at 72h (**Fig. S4c**). Loss of Tet2 and Tet3 almost eliminated the induced 5hmC modification at both enhancers and resulted in diminished expression of both *Aicda* mRNA and AID protein (**Fig. 4a, 4b**, **5b**; **Fig. S4c**), suggesting that TET proteins and 5hmC are required for the *TetE1* and *TetE2* enhancers to function at full capacity.

Studies from our lab and others have implicated TET proteins and 5hmC in regulating chromatin accessibility. For instance, TET proteins were shown to be required for demethylation of evolutionarily conserved enhancers during zebrafish development, and morpholino-mediated knockdown of *Tet1/2/3* resulted in decreased enhancer accessibility (*47*). In mammals, we have shown that *Tet2/3*-deficiency results in lower accessibility of enhancers during T and B cell development (*4, 34*). However, these steady state studies provide limited mechanistic insights. Here, through systematic analyses of 5hmC modification and chromatin accessibility kinetics during B cell activation, we show that 5hmC displays a time-dependent increase at regions that are differentially accessible between WT and *Tet2/3* DKO B cells during; moreover TET proteins are important for sustaining enhancer accessibility (**Fig. 6e; Fig. S6e**). We speculate that enhancer methylation limits enhancer output through recruitment of repressive complexes associated with a variety of proteins that bind methylcytosine or methylated CpGs (*48*), and that TET-mediated CpG hydroxymethylation and subsequent DNA demethylation are required to maintain enhancer accessibility, perhaps through recruitment of CXXC domain proteins such as Cpf1, a component of the SETD1 H3K4 methyltransferase complex (*49*).

How the activity of TET proteins is directed to specific sites in the genome remains an open question. In a previous study, we showed that TET proteins are recruited to Ig light chain enhancers during B cell development, potentially via the transcription factors PU.1 and E2A, and documented direct protein-protein interactions of Tet2 with both E2A and PU.1 (*4*). Here we propose that promiscuous interaction with transcription factors is a general mechanism for recruitment of TET proteins to specific loci, similar to the known interactions of p300 with hundreds of transcription factors (*50*). To identify the transcription factors mediating TET recruitment in activated B cells, we took advantage of our genome-wide analyses to assess enrichment for consensus transcription factor binding motifs in multiple datasets (5hmC, ATAC and DNA methylation). We found a consistent enrichment for NFκB, bZIP and bZIP:IRF composite binding motifs (Fig.s 1f, 2f; Fig. S1g, 6e). Based on a recent study showing that AP-1 (bZIP) family proteins, together with lineage-determining transcription factors, are “pioneer” factors capable of binding nucleosomal DNA and recruiting the BAF (SWI/SNF) complex for nucleosome remodeling (*51*), we focused on transcription factors of the bZIP family.

Our data suggest strongly that the bZIP transcription factor Batf is one of the major bZIP transcription factors responsible for TET recruitment to the *Aicda* locus. Batf is strongly induced at the mRNA level prior to *Aicda* induction in activated B cells (**Fig. 7a**), and Batf deficiency in B cells is associated with a dramatic impairment of AID expression and CSR (*40, 41*). Although loss of Tet2 and Tet3 had no significant effect on global Batf binding (**Fig. 7b; Fig. S7d**), Batf was required for 5hmC modification at *TetE1* (**Fig. 7e**). Composite bZIP:IRF motifs and AP-1 motifs that support BATF:JUN:IRF and BATF:JUN cooperation respectively were enriched in our genome-wide 5hmC, ATAC and DNA methylation datasets (**Fig. 1f**, **2f**; **Fig. S1g**, **6e**), consistent with previous findings that B cells lacking BATF, and IRF4/IRF8 show impaired *Aicda* induction and CSR (*40, 41, 52–54*). We propose that together, BATF:JunB and BATF:IRF assemblies recruit TET proteins as well as chromatin remodelling complexes to diverse enhancers including the *Aicda* enhancers *TetE1* and *TetE2* in activated B cells, thereby promoting enhancer accessibility, 5hmC deposition and DNA demethylation.

Our data emphasize the utility of 5hmC mapping for easy, one-step analysis of transcriptional and epigenetic landscapes in any cell type of interest. 5hmC is a quintessential epigenetic modification that marks the most highly enriched at active enhancers and the gene bodies of highly transcribed genes, and the relative levels of 5hmC at enhancers and gene bodies provide good estimates of enhancer function and the magnitude of transcription respectively (*46*). 5hmC mapping by CMS-IP sufficed to identify all known enhancers in the *Aicda* locus, in a manner that was superior to both H3K27Ac and Tet2 ChIP-seq, and changes in 5hmC identified enhancers relevant to any particular process of cell activation or differentiation separately from all enhancers in the genome. Given its high chemical stability, the fact that its measurement requires only purified DNA, and the availability of methods for its sensitive and specific detection, 5hmC is an appealing epigenetic mark for studying gene regulation. Overall, 5hmC distribution contains information analogous to those from ATAC-seq and ChIP-seq for enhancer histone modifications, effectively providing a transcriptional history of any given cell type written in DNA. If the genome is akin to an encyclopedia, 5hmC highlights those entries most relevant to a particular biological process.

## Acknowledgements

We would like to thank you Dr. Uttiya Basu for providing the AID antibody; Dr. Kefei Yu providing the Aicda-KO CH12F3 cells; Dr. Paolo Casali and Dr. Hong Zan for discussion; Laura Hempleman for assisting animal experiments; Cheryl Kim, Lara Nosworthy, Denise Hinz, and Robin Simmons (LJI Flow Cytometry Core) for help with cell sorting; Jeremy Day and Nick Wlodychak (LJI Functional Genomics Center) for assistance with next generation sequencing. C.-W.J.L. was supported by a Cancer Research Institute Irvington Postdoctoral Fellowship. V.S. is supported by Leukemia and Lymphoma Society Postdoctoral Fellowship. A.R. is supported by the National Institutes of Health (NIH) grants R35 CA210043 and R01 AI109842. D.G.S. supported by NIH Grant R01 AI127642. F.A. and A.C. have been partially supported by Institute Leadership Funds from La Jolla Institute for Allergy and Immunology and by the NIH Grant R01 MH111267. Funding for Illumina HiSeq 2500 and BD FACSAria II is supported by NIH (NIH S10OD016262, NIH S10RR027366).

## Author Contributions

C.-W.J.L. and V.S. conceptualized the experiments, acquired, analyzed the data, and wrote the manuscript. D.S.C. and E.G.A. performed majority of bioinformatics analysis (ATAC-seq, CMS-IP, enhancers analysis, WGBS, oxBS-seq, ChIP-seq), and proofread the manuscript. A.C. performed and advised on bioinformatics analysis (RNA-seq, TC-seq). C.-W.J.L. performed bioinformatics analysis (motifs, region characterization, data visualization) and the initial exploratory analysis. X.Y. provided key reagents and assistance for experiment (oxBS-seq). F.A. supervised the bioinformatics analysis and reviewed the manuscript. D.G.S. provided initial advised, key reagents, data interpretation, and reviewed the manuscript. A.R. supervised, conceptualized the experiments, data interpretation, and wrote the manuscript.

## Supplementary Tables

Supplementary Table S1.

Differentially expressed genes between WT and Tet2/3-DKO B cells, related to Fig. S4.

Supplementary Table S2.

Time-series analysis of RNA expression in WT B cells, related to Fig. S7.

Supplementary Table S3. Primers sequences.

Supplementary Table S4. Reagents and resource.

---

**Figure S1.**
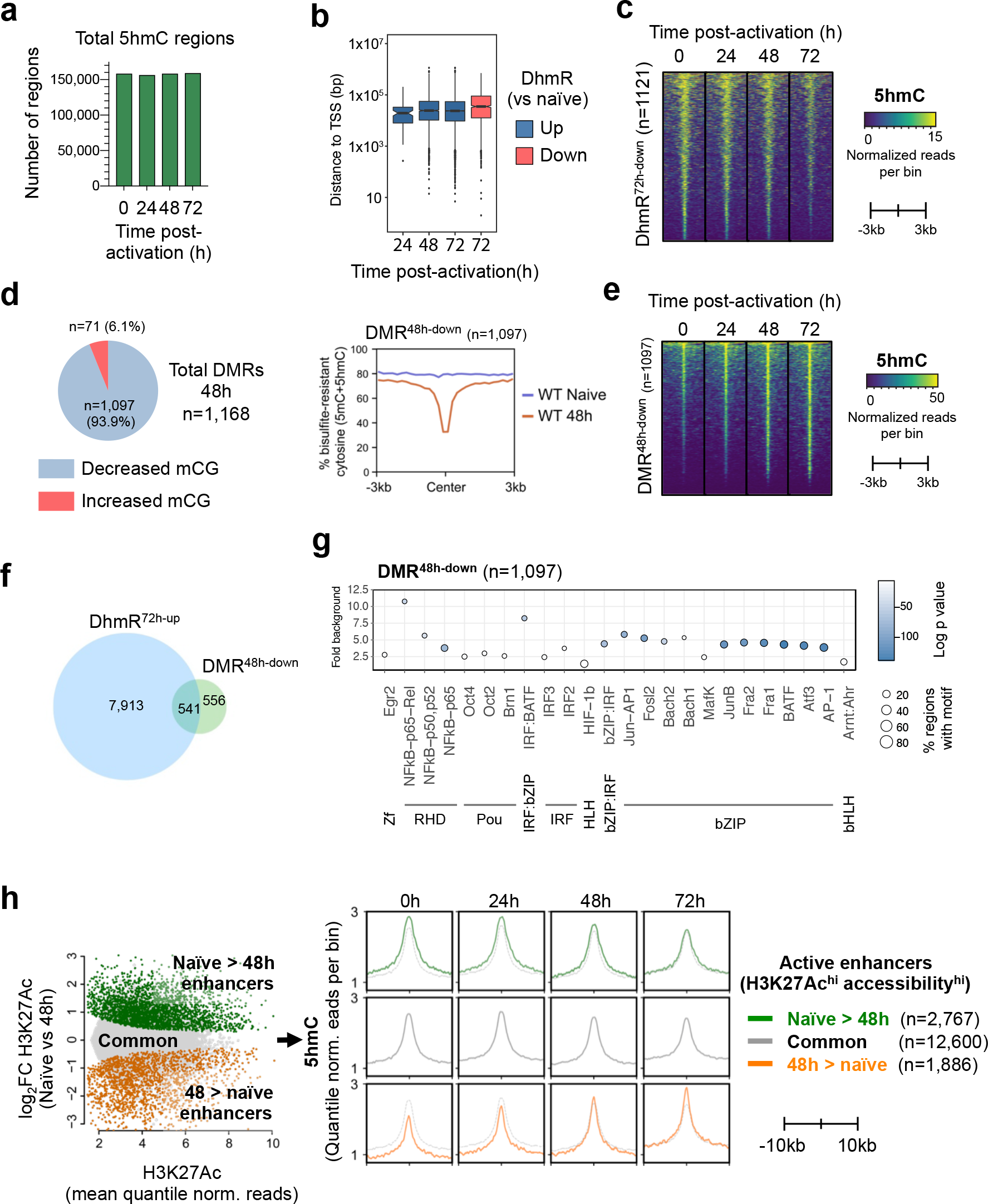
related to Fig. 1. Tet-mediated DNA hydroxymethylation correlates with demethylation and enhancer activity. **(a)** Similar total numbers of 5hmC-enriched regions between naïve and activated B cells. **(b)** Box-and-whisker plot showing the differential hydroxymethylated regions (DhmRs) in activated vs naïve B cells (see **Fig. 1c**) are located on average more than 10 kb from the closest transcription start site (TSS). **(c)** Heatmaps showing the kinetics of 5hmC modification at the 1,121 regions with decreased 5hmC in 72h-activated vs naïve B cells (DhmR^72h-down^; see **Fig. 1c**). 5hmC enrichment is shown as normalized reads per 100bp bin. **(d)** *Left*, the vast majority of differentially methylated regions (DMRs) with altered WGBS signal (5mC+5hmC) in naïve vs 48h-activated B cells show decreased DNA methylation. *Right*, plot of average DNA methylation (bisulfite-resistant cytosine 5mC+5hmC) at the DMR^48h-down^ (n=1,097) in 48h-activated vs naïve B cells. Average methylation is measured per 200 bp bin. **(e)** Heatmaps showing the kinetics of 5hmC at the 1,097 DMR^48h-down^ with decreased methylation 48h post-activation. 5hmC enrichment is shown in normalized reads per 100 bp bin. **(f)** Venn diagram showing the overlap between regions with demethylation (DMR^48h-down^) and hydroxymethylation (DhmR^72h-up^). Note that 49.3% (541/1097) of DMR are progressively hydroxymethylated post-activation. **(g)** Strong enrichment for consensus binding motifs of NFκB, IRF:bZIP, bZIP, and other transcription factors in the 1,097 demethylating regions after 48h activation (DMR^48h-down^). Random genomic regions were used as background for motif analysis. Y-axis indicates the fold enrichment versus background, circle size indicates the percentage of regions containing the respective motif, and the color indicates the significance (Log_10_ *p* vaule). **(h)** 5hmC level tracks with enhancer activity. *Left*, differential active enhancers between naïve and 48h-activated B cells were classified based on the significant difference in H3K27Ac and accessibility (ATAC-seq) into preferential naïve-B-active enhancers (green, “Naïve>48h”) and activated-B-active enhancers (orange, “48>naïve”). The remaining enhancers not meeting the above criteria were classified as common active enhancers (grey, “Common”). *Right*, mean level of 5hmC per bin (50 bp) at the +/− 10kb interval to the center was plotted for each type of active enhancers. Note that the 5hmC levels from the “Common” enhancers (*middle)* are also plotted as dotted lines for naïve-B-active (*top*) and activated-B-active enhancers (*bottom*) as reference.

**Figure S2.**
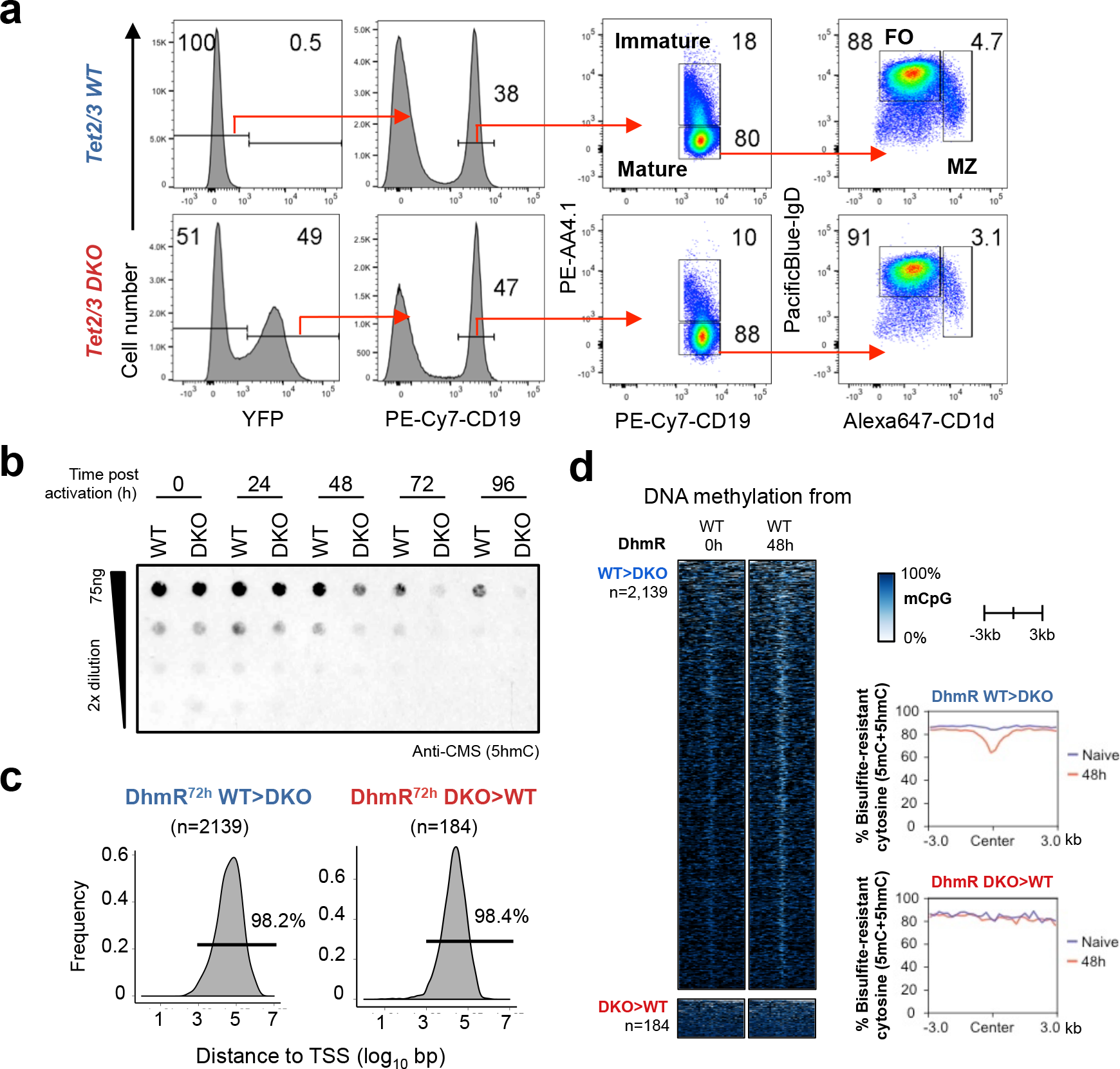
related to Fig. 2. Phenotypic features of WT and *Tet2/3* DKO B cells. **(a)** Comparable splenic mature B cell populations in *Tet2/3*-conditional deletion mice. WT (*Tet2/3-flox Rosa26-LSL-YFP*) and DKO (*Cre^ERT2^ Tet2/3-flox Rosa26-LSL-YFP*) mice were treated as in **Fig. 2b** and the phenotype of splenic B cells was analyzed on day 7 after the initial tamoxifen injection. Plots were first gated on live single cells based on FSC/SSC (*first panel*) and total (WT) or YFP^+^ (DKO) CD19^+^ B cells were subsequently gated (*second panel*), followed by the analysis of mature and immature B cells (*third panel*); and follicular (FO) and marginal zone (MZ) B cells (fourth panel). **(b)** Total 5hmC levels in WT and *Tet2/3* DKO B cells assessed by cytosine 5-methylenesulphonate (CMS) dot blot (see *Materials and Methods*). Note that 5hmC levels decrease in *Tet2/3* DKO B cells only after several rounds of cell division (>48h). **(c)** Histograms showing the distance from the TSS to the TET-regulated DhmR regions differentially marked with 5hmC in 72h-activated *Tet2/3* DKO relative to WT B cells (see **Fig. 2d, 2e**). The 2,139 and 184 DhmRs with decreased (*left*, DhmR^72h^ WT>DKO) and increased (*right*, DhmR^72h^ DKO>WT) 5hmC after activated for 72h 72h-activated *Tet2/3* DKO relative to WT B cells are located on average more than 10 kb from the closest TSS. **(d) Tet-mediated 5hmC modifications mark DNA demethylation**. *Left panels*, heatmaps show the DNA modification status from naïve and 48h-activated WT B cells (WGBS, 5mC+5hmC) at the 2,139 and 184 DhmR regions with decreased (*top*, WT>DKO) and increased (*bottom*, DKO>WT) 5hmC in 72h-activated *Tet2/3* DKO vs WT B cells. *Right panels*, plots of the average decrease in bisulfite-resistant modifications (5mC+5hmC) per bin (200 bp) at these regions. The majority of the 2,139 WT>DKO DhmRs at 72h show decreased modifications in activated WT B cells (*top*); the 184 DKO>WT DhmR at 72h are fully modified and unchanged in *Tet2/3* DKO vs WT (*bottom*).

**Figure S3.**
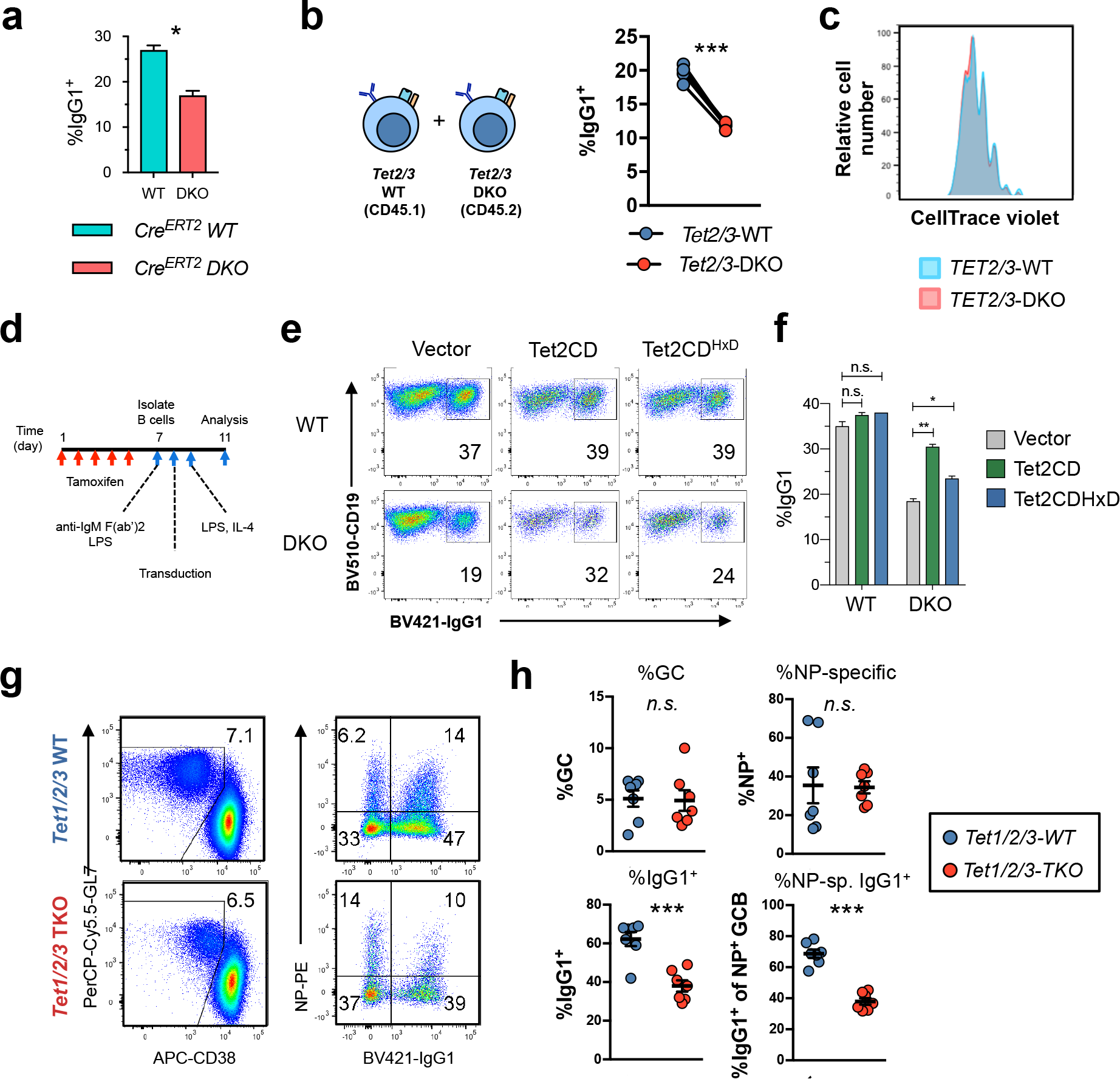
related to Fig. 3. TET family proteins are important for B-cell-intrinsic CSR. **(a) CSR defect is not caused by Cre activity**. *Cre^ERT2^* Rosa26-LSL-YFP (*Cre^ERT2^ WT*) and *Cre^ERT2^ Tet2^fl/fl^ Tet3^fl/fl^ Rosa26-LSL-YFP* (*Cre^ERT2^* DKO) mice were injected with tamoxifen as in **Fig. 2b**. Isolated B cells were activated with LPS and IL-4 in the presence of 4-hydroxytamoxifen for 4 days and %IgG1^+^ cells were analyzed (gated on live CD19^+^ YFP^+^). One experiment is shown. n=2 for each genotype. **(b-c) (b) CSR defect is cell-intrinsic. (b)** *(Left) Tet2^+/+^Tet3^+/+^* WT CD45.1 and *Tet2/3-*DKO CD45.2 mice were treated as in **Fig. 3a** and isolated B cells were labeled with Cell-Trace violet, 1:1 mixed, and activated with LPS and IL-4 for 4 days. *(Right)* Cells were gated based on CD45.1 and CD45.2 and the percentages of IgG1-switched cells in WT and DKO are shown. Cells from the same well are connected with lines. **(c)** Co-cultured WT and *Tet2/3*-DKO B cells showed similar proliferation profiles. Data shown are representative of two independent experiments with four technical replicates for each genotype. **(d-f)(d)** Flow chart of experiment to assess the importance of TET catalytic activity in CSR. **(e)**Flow cytometry plots and **(f)** quantification of WT and *Tet2/3* DKO B cells transduced with empty vector (*left panels*), Tet2 wild-type catalytic domain (Tet2CD, *middle panels*), and Tet2 HxD mutant catalytic domain (Tet2CD^HxD^, *right panels*) shows that TET catalytic activity can partly rescue the CSR to IgG1. Data shown are representative of two independent experiments with two technical replicates. n.s., not significant. ***, *p<0.01*. *, *p<0.05*. **(g-h) Deletion of all TET proteins Tet1/2/3 showed similar decrease in CSR compared to** *Tet2* **/3-DKO**. **(g)** *Tet1/2/3-flox Cre^ERT2^ Rosa26-LSL-YFP* (TKO) and control *Tet1/2/3-flox Rosa26-LSL-YFP* (WT) mice were treated with tamoxifen and immunized with NP-OVA as in **Fig. 3a**and GC response and CSR were analyzed on day 7. Flow cytometry plots showed the percentage of GCB (CD38^lo^GL7^hi^) in WT and *Tet1/2/*3-*TKO* lymph node cells gated on total (WT) and YFP^+^ (TKO) live CD19^+^ B cells (*left panels)*. Antigen-specific (NP-PE) and class-switched cells (IgG1) were analyzed among the GCB. **(h)**Quantification of the experiments showed in **(g)**. Data shown are aggregated results from two independent experiments. WT, n=7; TKO, n=7. Statistical significance was calculated using unpaired two-tailed *t*-test. n.s., not significant. ***, *p<0.01*. *, *p < 0.05*.

**Figure S4.**
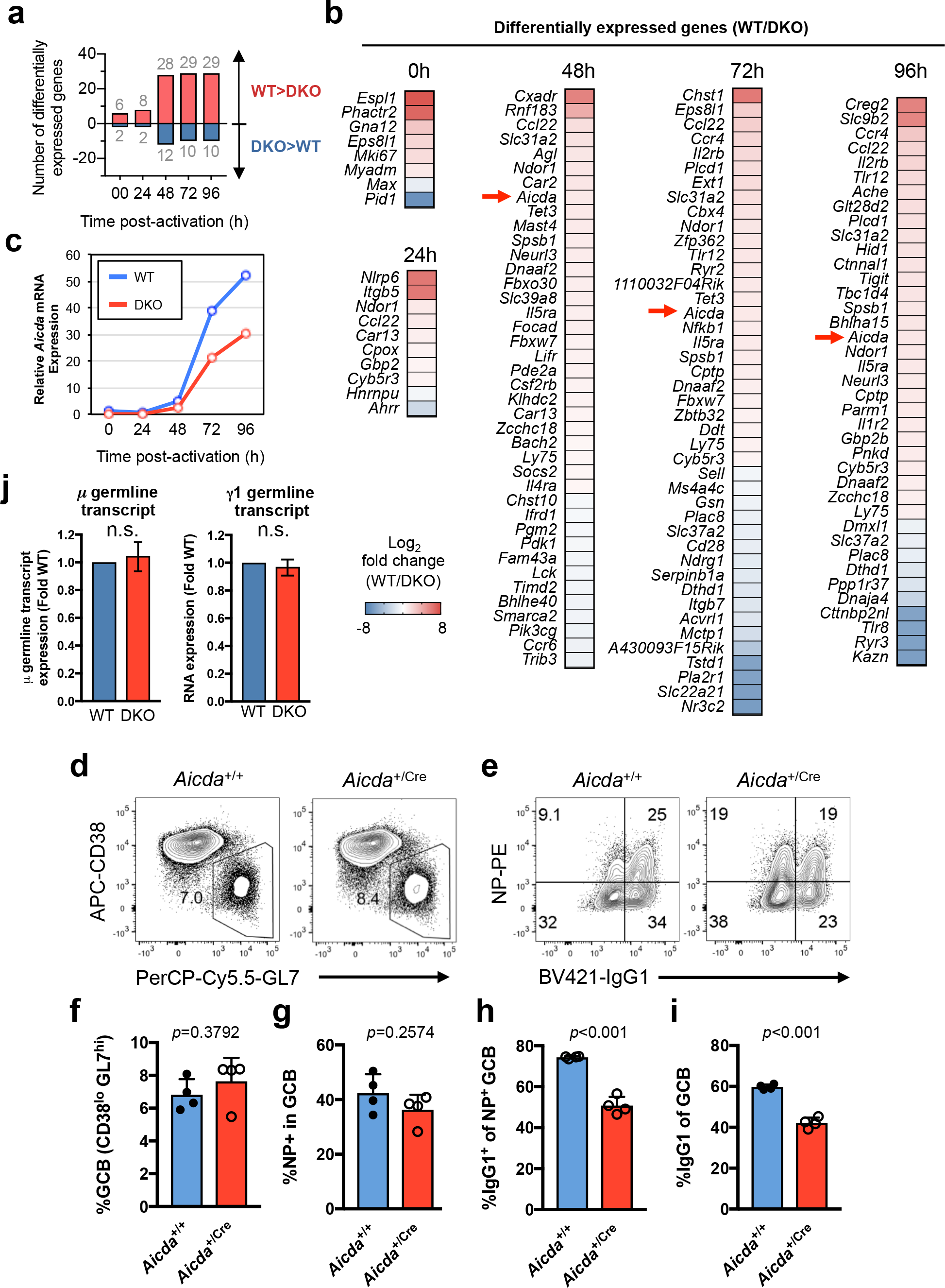
related to Fig. 4. Decreased *Aicda* expression in*Tet2/3-*DKO B cells. WT and *Tet2/3-* DKO B cells were activated as in **Fig. 3g** and the transcriptomes were analyzed by RNA-seq (see *Materials and Methods* and **Supplementary Table S1** for details). **(a)** Number of differentially expressed genes between WT and *Tet2/3*-DKO B cells as a function of time after activation. Relatively few genes show alterations in their expression. **(b)** List of all differentially expressed genes between WT and *Tet2/3*-DKO B cells. *Aicda* (indicated by red arrows) was one of the genes expressed significantly lower in DKO. Color scale indicates Log_2_ fold change between WT and DKO. **(c) Tet2 and Tet3 are required for potent *Aicda* expression**. *Aicda* mRNA expression was analyzed by qRT-PCR as in **Fig. 4a**as a function of time after activation. Results showed a consistent 50% decrease in *Tet2/3* DKO relative to WT B cells. **(d-i) Haploinsufficiency of *Aicda* results in decreased CSR**. Mice with indicated genotypes were immunized with 10ug of NP-OVA mixed with Alum via footpad and the draining lymph nodes were analyzed by FACS day 7 post-immunization. Heterozygous *Aicda-Cre* mice were used to model *Aicda* haploinsufficiency as the knocked-in Cre recombinase disrupted the endogenous *Aicda* expression. Representative FACS analysis of **(d)** germinal center B cells (GCB; CD38^lo^ GL7^hi^) and **(e)** CSR to IgG1. **(f-i)** Statistical analyses of the populations are shown (n=4 each). Data shown are representative of two independent experiments. Means and standard errors are shown. Unpaired two-tailed *t*-test was used to calculate statistical significance and the *p* values are indicated. **(j) Tet2 and Tet3 are not required for expression of germline transcripts**. WT and DKO B cells were activated for 4 days, and *μ*and *γ*1 germline transcripts were analyzed by qRT-PCR. Data were normalized to *Gapdh* and to WT level as in **Fig. 4a**. n.s., not significant.

**Figure S5.**
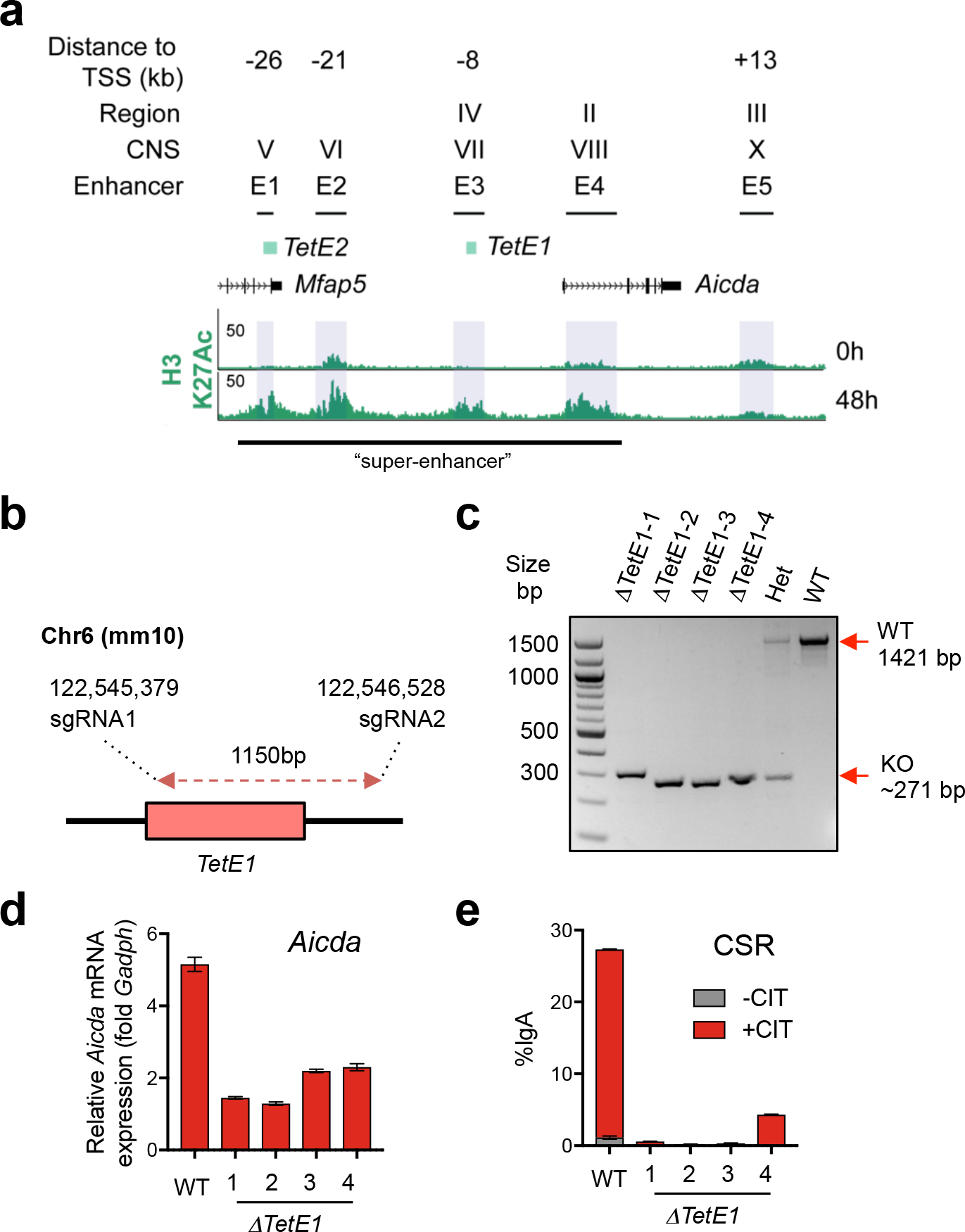
related to Fig. 5. The TET-responsive element*TetE1* regulates CSR and*Aicda* mRNA expression in the CH12 B cell line. **(a)** Diagram depicts the relative position of TET-responsive elements *TetE1* and *TetE2* to previously identified *Aicda* distal and intronic enhancers. “Region” IV, II, III are from Tran et al., 2010; “CNS” V-X are from Crouch et al., 2007; “Enhancer” E1-E5 from Kieffer-Kwon et al., 2013. Note that the promoter-proximal element is not depicted. Coordinate for the shown locus is chr6:122,523,500-122,576,500 (mm10). **(b-e) *TetE1* is important for regulating*Aicda* expression and CSR. (b)** Scheme for *TetE1* deletion in CH12 cells with CRISPR. **(c)** Four clones were identified with homozygous deletion of *TetE1* as examined by PCR followed by gel electrophoresis. A clone with heterozygous deletion (Het) and a WT control were showed as controls. **(d-e)** WT and TetE1-deletion clones were stimulated with CIT (anti-CD40, IL-4, TGFβ) for two days. **(d)** *Aicda* mRNA expression and **(e)** CSR to IgA was analyzed by qRT-PCR and flow cytometry, respectively. Results showed that deletion of *TetE1* decreased both *Aicda* mRNA expression and abrogated CSR.

**Supplementary Figure S6.**
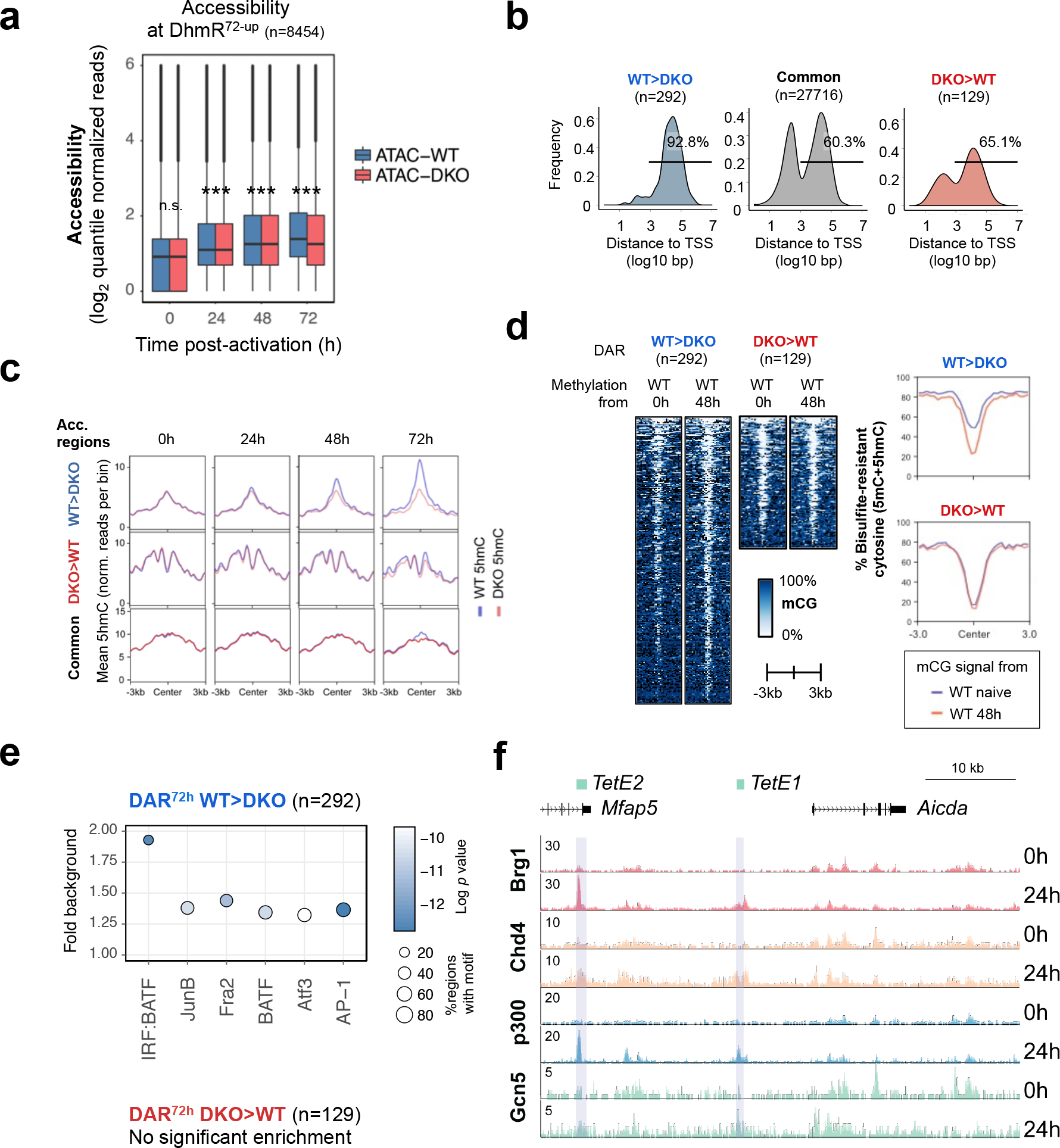
related to Figure 6. Analysis of Tet-dependent accessible regions. **(a) Correlation between 5hmC and accessibility**. At the differentially 5hmC-enriched regions between WT and DKO (DhmR^72h-up^, n=8454, **Fig. 1c**), mean chromatin accessibility detected by ATAC-seq was shown for each region. Note that the closely correlated increase of both 5hmC and accessibility (blue, ATAC-WT). Loss of Tet2/3 resulted in an observed decrease in accessibility, corresponding to the results from **Fig. 6c**. Statistical significance between WT and DKO at each time point was calculated by Kolmogorov-Smirnov test with Bonferroni correction using the family-wise error rate. n.s., not significant. ***, p adj. < 0.01. The exact adjusted *p* values are 0.06, 6.01e-05, 1.79e-07, 3.611e-11 for 0h, 24h, 48h, 72h, respectively. **(b) Tet-facilitated accessibility at distal elements**. Histograms showing the distance of DARs (WT>DKO and DKO>WT) and commonly accessible regions (Common) from the closest TSS. Majority of the Tet-facilitated accessible regions (WT>DKO, n=292) are distal elements (>1000bp; 92.8%). **(c) Tet2/3-dependent accessible regions are hydroxymethylated**. Line plots showing the kinetics of mean 5hmC modification at the differential accessible regions (DARs) between WT and Tet2/3-DKO for the data depicted in **Fig. 6c**. Regions that are more accessible in WT (WT>DKO, n=292), less accessible in WT (DKO>WT, n=129), and commonly accessible (n=27,716) are shown on top, middle, and bottom panels, respectively. 5hmC enrichment is shown as normalized reads per 100 bp bin. Note the Tet-dependent 5hmC modification at these DARs (*compare 72h panels*). **(d) Tet-facilitated accessible regions further demethylated after activation**. *Left,* heatmaps showing the DNA modifications status (5mC+5hmC) in naïve and 48h-stimualted WT B cells at the WT>DKO DARs or Tet-facilitated accessible regions (n=292) and DKO>WT DARs (n=129). *Right*, plots summarized the results for the heatmaps (*left*) with y-axis indicates the level of bisulfite-resistant cytosine (5mC+5hmC). In WT B cells, regions that lose accessibility in *Tet2/3* DKO B cells relative to WT (WT>DKO) also show a decrease in modifications (mostly 5mC) after activation. **(e)** Enrichment for consensus IRF:bZIP (IRF:BATF) and bZIP transcription factor binding motifs in the 292 Tet-facilitated accessible regions, which shows increased accessibility in WT relative to *Tet2/3* DKO B cells at 72h. No significant motif enrichment was detected at the DKO>WT DAR (n=129). Commonly accessible regions were used as background for the analysis. Y-axis indicates the fold enrichment versus background, circle size indicates the percentage of regions containing the respective motif, and the color indicates the significance (Log_10_ *p* vaule). **(f) B cell activation induces recruitment of chromatin regulators to** *Aicda* **distal elements**. Genome browser view of ChIP-seq data showing B cell activation induces the binding of the chromatin remodelling complex components Brg1 and Chd4, and histone acetyltransferases p300 and Gcn5 to the TET-responsive *Aicda* elements *TetE1* and *TetE2* in naïve and activated WT B cells. Scale indicates reads per 10 millions. Coordinate for locus is chr6:122,523,500-122,576,500 (mm10).

**Figure S7.**
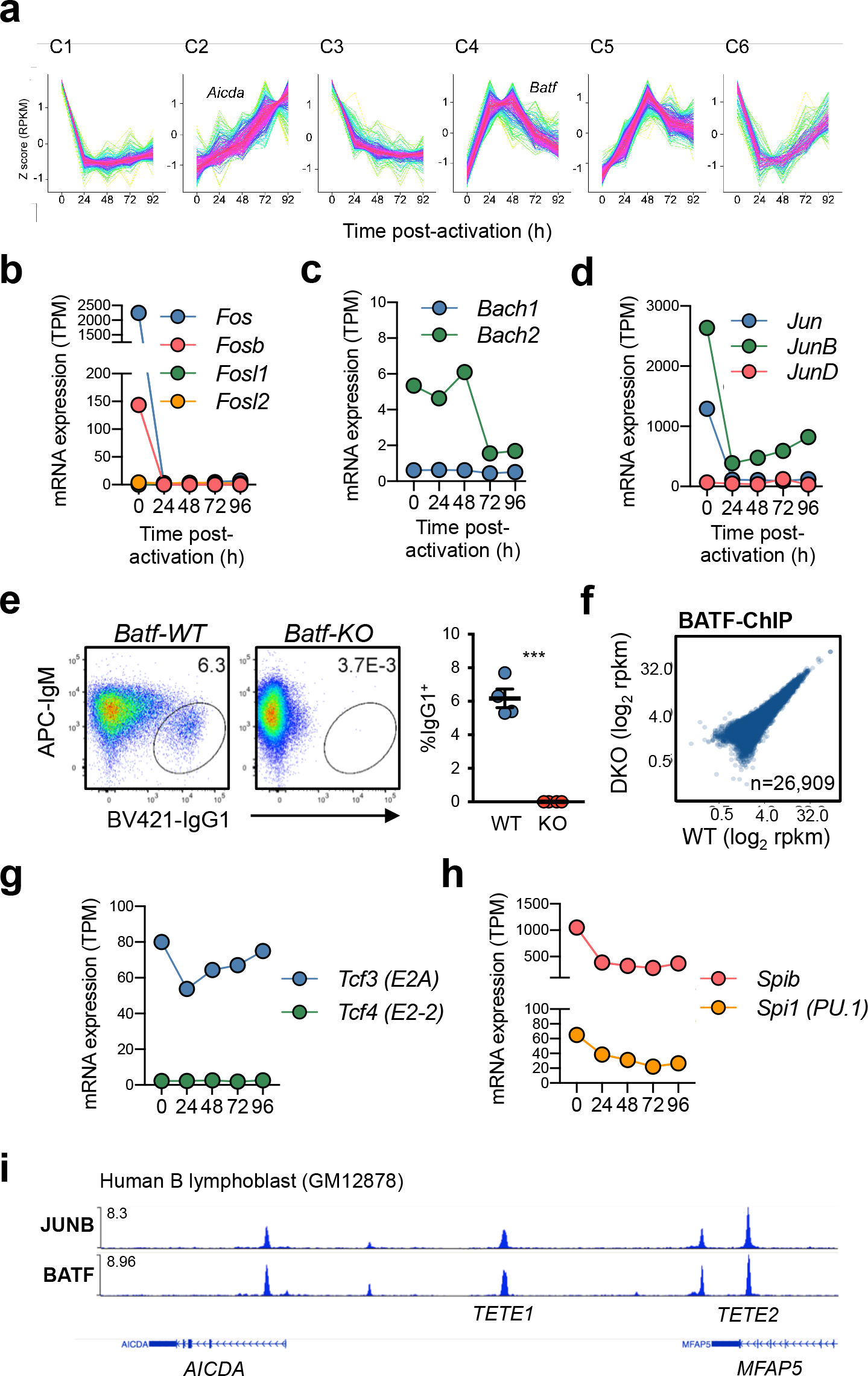
related to Fig. 7. AP-1 proteins in activated B cells. **(a) Analysis of temporal gene expression modules.** Gene expression in WT cells activated for various times were analyzed and genes were clustered based on the temporal expression patterns. Six clusters (C1-C6), or expression patterns, were identified. *Aicda* and *Batf* are found in C2 and C4, respectively. Y-axis indicates the Z-score calculated using RPKM. X-axis indicates the time post-activation. For details, see *Materials and Methods* and **Supplementary Table S2**. **(b-d) Expression of AP-1 proteins**. Mean mRNA expression of selected AP-1 proteins in naïve and activated B cells including *Fos* family **(b)**, *Jun* family **(c)**, and *Bach* family **(d)**. Note that LPS/IL-4 simulation decreased the expression of *Fos, Fosb, Jun,* and *JunB* followed by a gradual increase in *JunB* expression (**b, c**). The high basal expression of these factors could potentially attribute to minor B cell subset, including memory B cells. **(d)** Expression of *Bach2,* a transcription factor expressed in naïve and memory cells, also decreased with a delayed kinetics at 72h post-activation. Data shown are from RNA-seq with two independent replicates. TPM, transcript per million. **(e)** Batf is required for CSR. B cells were isolated from WT and *Batf-KO* and activated with LPS and IL-4 for 4 days. Class switch recombination to IgG1 was analyzed by flow cytometry (gated on live CD19^+^). Data shown is representative of two independent experiments (n=4 each). Means and standard errors are shown. Statistical significance was calculated using unpaired two-tailed *t*-test. ***, *p*<0.01. **(f) Tet proteins are not necessary for genome-wide BATF binding**. Plots show the highly similar distribution of BATF in WT and DKO 72h-activated B cells as analyzed by ChIP-seq with two independent replicates. Data shown are the comparison of the BATF enrichment in WT and DKO at the 26,909 regions integrated from the joined peaks from two replicates each of WT and DKO. Axes depict the log_2_ rpkm (read per kilo base per million) using quantile-normalized reads for each regions analyzed. Note that no region was significantly differenct between WT and DKO using an adjusted *p* value at 0.05. **(g-h) Expression of E-box and Ets family proteins**. Mean mRNA expression for E-box and Ets family proteins are shown. Data shown are from RNA-seq with two independent replicates. TPM, transcript per million. **(i) JUNB and BATF bind to** *AICDA* **enhancers in human B cells**. JUNB and BATF bindings in human B cell lymphoblast GM12878 at the *AICDA* locus are shown (Hg38; chr12:8,598,356-8,655,770). The approximate locations for *TETE1* and *TETE2* are indicated. Data were originally from ENCODE project, processed by CistromeDB, and were viewed using WashU Epigenome Browser.

## Materials and Methods

### Mice

*Tet2*^*fl/fl*^ and *Tet3*^*fl/fl*^ mice were generated as previously described^55, 56^. C57BL/6J (000664), Ubc-CreERT2 (008085; described as Cre^ERT2^ herein), Rosa26-LSL-EYFP (006148), and AID-Cre (007770) were obtained form Jackson Laboratory. All mice used were 8-16 weeks in the C57BL/6 background and kept in specific-pathogen free animal facilitate at La Jolla Institute and were used according to protocols approved by the Institutional Animal Care and Use Committee. To induce Cre^ERT2^-mediated deletion, Cre-expressing and control mice were intraperitoneally injected with 2 mg tamoxifen (Sigma) dissolved in 100 uL corn oil (Sigma) daily for 5 days.

### Immunization

For 4-hydroxy-3-nitrophenylacetyl-conjugated ovalbumin (NP-OVA; Biosearch) immunization, the hapten-conjugated protein was diluted to 1 mg/mL in PBS was mixed with 1 volume of Alhydrogel (Invivogen) and injected into hind footpads (10 ug in 20 uL per injection). Germinal center response was analyzed 7 days later and the two draining popliteal lymph nodes were pooled for analysis. Hapten-specific B cells were identified by positive staining with NP-phycoerythrin (BioSearch Technologies).

### B cell isolation and class switch recombination (CSR)

B cells were isolated with EasySep Mouse B cell isolation kit (Stem Cell Technology, Canada) from splenocytes. To induce class switch recombination from IgM to IgG1, B cells (5×10^5^-1×10^6^ cells/ mL) were activated with 25 ug/mL LPS from *E. coli* O55:B5 (Sigma, St. Louis, MO) and 10 ng/mL rmIL-4 at 37°C 5% CO_2_. For IgA switching, cells were activated with anti-CD40 (1 ug/mL, clone 1C10, Biolegend), rmIL-4 (10 ng/mL, Peprotech), rmIL-5 (10 g/mL, Peprotech), and rhTGFβ1 (1 ng/mL). Media were composed of RPMI1640 (Thermo Fisher, Waltham, MA) supplemented with 10% FBS, 1× MEM non-essential amino acids, 10 mM HEPES, 2 mM Glutamax, 1 mM sodium pyruvate, 55 uM 2-mercaptoethanol, and 50 ug/mL gentamicin (all from Thermo Fisher, Waltham, MA). To enhance *Cre*^*ERT2*^-mediated deletion, cells from *Cre*^*ERT2*^ mice were cultured in the presence of 1 uM 4-hydroxytamoxifen (Tocris). All cytokines used above were from Peprotech (Rocky Hill, NJ).

### Retroviral transduction and two-step CSR

Retrovirus was produced by transfecting PlatE cells with MSCV-based retroviral vectors and pCL-Eco. Naïve B cells were stimulated with 5 ug/mL F(ab’)_2_ goat anti-mouse IgM (Jackson Immuno Research) and 10 ug/mL LPS at 1×10^6^ cells/mL for 24-48 hours. Retrovirus was added to the cells in the presence of 20mM HEPES and 0.8 ug/mL Polybrene (Millipore) and centrifuged at 3,000 rpm at 32°C for 90 mins. Cells were transferred back to 37°C 5% CO_2_ incubator for another 24 hours. To induce CSR, cells were washed once with warm media and activated with LPS and IL-4 as above for 48 hours. Under this condition, CSR was inhibited and started to class switch only after LPS/IL-4 activation.

### Flow cytometry

Primary cells and *in vitro* cultured cells were stained in FACS buffer (1% bovine serum albumin, 1mM EDTA, and 0.05% sodium azide in PBS) with indicated antibodies for 30 mins on ice. Cells were washed and then fixed with 1% paraformaldehyde (diluted from 4% with PBS; Affymetrix) before FACS analysis using FACS Canto II and FACS LSR II (BD Biosciences). Antibodies and dye were from BioLegend, eBioscience, and BD Pharmingen. Data were analyzed with FlowJo (FlowJo LLC, Ashland, OR).

### Immunoblotting

Proteins isolated from B cells with RIPA buffer were resolved using NuPAGE 4-12% Bis-Tris gel (ThermoFisher) and transferred from gel to PVDF membrane using Wet/Tank Blotting Systems (Bio-Rad). Membrane was blocked with 5% non-fat milk (Bob’s red mill) in TBSTE buffer (50 mM Tris-HCl pH 7.4, 150 mM NaCl, 0.05% Tween-20, 1 mM EDTA), incubated with indicated primary antibodies, followed by secondary antibodies conjugated with horseradish peroxidase (HRP) and signal was detected with enhance chemiluminescence reagents and X-ray film.

### RNA extraction, cDNA synthesis, and quantitative RT-PCR

Total RNA was isolated with RNeasy plus kit (Qiagen, Germnay) or with Trizol (ThermoFisher, Waltham, MA) following manufactures’ instruction. cDNA was synthesized using SuperScript III reverse transcriptase (ThermoFisher) and quantitative RT-PCR was performed using FastStart Universal SYBR Green Master mix (Roche, Germany) on a StepOnePlus real-time PCR system (ThermoFisher). Gene expression was normalized to *Gapdh*. Primer are listed in **Supplementary Table 3**.

### Bisulfite- (BS) and oxidative-bisulfite-(oxBS) sequencing

The BS and oxBS procedures were performed as previously described^28^. Briefly, three PCR products containing C, mC, or hmC pertaining to different regions of λ phage genome were used as spike-ins at a ratio of 1:200 of the genomic DNA. 1.5 µg of genomic DNA mixed with spike-ins was ethanol precipitated of which 1 µg of the DNA was oxidized with potassium perruthenate (KRuO_4_; Sigma) prior to bisulfite (BS) treatment (for oxBS) using MethylCode bisulfite conversion kit (ThermoFisher) and 0.5 µg of DNA was directly used for BS treatment. The BS and oxBS treated DNA were amplified using respective PCR primers and as well as primers specific to the spike-in PCR products with KAPA Uracil^+^ PCR mix (Roche). The amplified products were pooled and libraries were prepared using the NEB Ultra II library preparation kit (NEB) according the manufacturer. The libraries were sequenced paired-end 250bp by 250bp using MiSeq with the MiSeq reagent kit v2 (500-cycles; Illumina). Primer are listed in **Supplementary Table 3**.

### Genome-wide 5hmC mapping by cytosine-5-methylenesulfonate immunoprecipitation (CMS-IP)

CMS-IP was performed similar to perviously described^4, 24, 25^. Briefly, genomic DNA isolated from naïve and activated B cells was spiked with unmethylated lambda phage cI857 Sam7 DNA (Promega, Madison, WI) and a PCR amplicon from puromycin resistant gene at a ratio of 200:1 and 100,000:1, respectively. DNA (5-10 ug in 130 uL TE buffer) was sheared with a Covaris E220 (Covaris) using microTUBE for 4 mins. DNA was cleaned-up with AmureXP beads, processed with NEBNext End Repair and A-tail Modules (NEB, Ipswich, MA), and ligated to methylated Illumina adaptors (NEB). DNA was then bisulfite-treated (MethylCode, ThermoFisher), denatured, and immunoprecipitated with anti-CMS serum (in house) and mixture of protein A and G dynabeads (ThermoFisher). Libraries for immunoprecipitated DNA were generated by PCR with barcoded primers (NEBNext Multiplex Oligos for Illumina, NEB) for 15 cycles using KAPA HiFi HotStart Uracil+ ReadyMix (Roche), followed by a cleanup with AmpureXP beads (Beckman Coulter), and sequenced with a HiSeq2500 (Illuminia, San Diego, CA) with paired-end 50 bp reads.

### Locus specific analysis of 5hmC with AbaSI-qPCR

Genomic DNA (200 ng) was treated with T4 beta-glucosyltransferase (ThermoFisher) in the presence of UDP-glucose to glycosylate 5hmC at 37°C overnight. Half of the reaction was digested with AbaSI (NEB), which is specifically active for glycosylated 5hmC, for 4 hours at 25°C followed by 15 mins at 65°C to inactivate enzymes. The uncut sample was processed as above without the addition of AbaSI. Equal amount of DNA from the above reactions was used as template for real-time PCR as described for RNA qRT-PCR using primers TetE1-CMS-qF and TetE1-CMS-qR. To monitor the degree of digestion, samples were spiked-in 1 pg control DNA with a single 5hmC-modified CpG (EpiMark 5-hmC and 5-mC Analysis Kit; NEB). The relative amount of 5hmC was calculated by the percentage of decrease in qPCR signals in digested half relative to undigested half. As control to monitor non-specific digestion (not show), a genomic region containing CpG motifs but without 5hmC modification in B cells (*Foxp3 CNS2*) was used as a control with Foxp3-CNS2-qF and Foxp3-CNS2-qR. Primers are listed in **Supplementary Table 3**.

### DNA dot blot

DNA dot blot was performed as previously described^4, 28^. To analyze 5hmC abundance, genomic DNA was treated with sodium bisulfite as above. DNA was diluted two-fold serially with TE buffer, denatured in 0.4 M sodium hydroxide and 10 mM EDTA at 95°C for 10 mins, and then immediately chilled on ice. Equally volume of ice-cold 2 M ammonium acetate pH 7.0 was added and incubated on ice for 10 mins. Denatured DNA were spotted on a nitrocellulose membrane using a Bio-Dot apparatus (Bio-Rad), washed with 2× SSC buffer (300 mM NaCl and 30 mM sodium citrate), and baked in a vacuum oven at 80°C for 2 hours. To detect CMS, membrane was rehydrated with TBSTE buffer and blocked with 5% non-fat milk (Bob’s red mill) in TBSTE buffer. CMS was detected with primary rabbit anti-CMS antisera (in house) following the procedures above for Immunoblotting.

### Chromatin Immunoprecipitation sequencing (ChIP-seq)

Chromatin immunoprecipitation was performed as described before^4^. Briefly, cells were fixed with 1% formaldehyde (ThermoFisher) at room temperature for 10 mins at 1×10^6^ cell/mL in media, quenched with 125 mM glycine, washed twice with ice cold PBS. Cells were pelleted, snap-froze with liquid nitrogen, and store at −80°C until use. For Tet2-ChIP, activated cells were centrifugated at 250 × *g* for 5 mins and cell pellets were resuspended in 37°C PBS with 2mM disuccinimidyl glutarate to crosslink proteins for 30 mins at room temperature. Formaldehyde was added to a final concentration of 1% and the cells were incubated at room temperature for 10 mins with nutation. Quenching and cell storage were performed as above. To isolate nuclei for sonication, cell pellets were thawed on ice and lysed with lysis buffer (50 mM HEPES pH 7.5, 140 mM NaCl, 1mM EDTA, 10% glycerol, 0.5% NP40, 0.25% Triton-X100) for 10 mins at 4°C with rotation, washed once with washing buffer (10 mM Tris-HCl pH 8.0, 200 mM NaCl, 1 mM EDTA, 0.5 mM EGTA) and twice with shearing buffer (10 mM Tris-HCl pH 8.0, 1 mM EDTA, 0.1% SDS). Nuclei were resuspended in 1mL shearing buffer and sonicated with Covaris E220 using 1 mL milliTUBE (Covaris, Woburn, MA) for 18-20 minutes (Duty Cycle 5%, intensity 140 Watts, cycles per burst 200). After sonication, insoluble debris was removed by centrifugation at 20,000 × *g*. Buffer for chromatin was adjusted with 1 volume of 2× conversion buffer (10 mM Tris-HCl pH 7.5, 280 mM NaCl, 1 mM EDTA, 1mM EGTA, 0.2% sodium deoxycholate, 0.2% Triton-X100, 1% Halt protease inhibitors with (for H3K27Ac) or without (for BATF, Tet2) 0.1% SDS. Chromatin was pre-cleared with washed protein A dynabeads (ThermoFisher) for 2 hours, incubated with antibodies and protein A dynabeads overnight (all procedures were at 4°C with rotation). For H3K27Ac ChIP, bead-bound chromatin was washed twice with RIPA buffer (50 mM Tris-HCl pH 8.0, 150 mM NaCl, 1 mM EDTA, 0.5% sodium deoxycholate, 1% NP-40, 0.1% SDS), once with high salt wash buffer (50 mM Tris-HCl pH 8.0, 500 mM NaCl, 1 mM EDTA, 1% NP-40, 0.1% SDS), and once with TE (10 mM Tris-HCl pH 8.0, 1 mM EDTA). For BATF ChIP, all wash buffers were as above but without SDS. For Tet2 ChIP, beads were washed three times with RIPA buffer without SDS and once with TE. Chromatin was eluted from beads with elution buffer (100 mM NaHCO_3_, 1% SDS, 1 mg/mL RNaseA; Qiagen) twice for 30 mins each at 37°C with constant shaking. NaCl and proteinase K (Ambion) were added to the eluted chromatin at concentrations of 250 mM and 0.5 mg/mL, respectively, and de-crosslinked at 65°C overnight with constant shaking. DNA was purified with Zymo ChIP DNA Clean & Concentrator-Capped Column (Zymo Research, Irvine, CA). Library was prepared with NEB Ultra II library prep kit (NEB) following manufacture’s instruction and was sequenced on an Illumina Hiseq 2500 (single-end 50 bp reads).

### ATAC-seq

Procedures were as described^4^. Briefly, 50,000 cells were collected by centrifugation and washed once with 50uL ice-cold PBS and centrifuged at 600 *xg* for 5 mins at 4°C. Cell pellets were resuspended in 50 uL of cold lysis buffer (10mM Tris-HCl pH 7.4, 10 mM NaCl, 3 mM MgCl_2_, 0.1% IGEPAL CA-630) and spin down immediately at 500 *xg* for 10 mins at 4°C. Supernatant was discarded and nuclei were resuspended in 50uL transposition reaction mix (25uL 1× TD buffer fom Illuminia, 2.5 uL Tn5 transposase, 22.5 uL H_2_O), incubated at 37°C for 30 mins, and DNA was purified with a Qiagen MinElute kit (Qiagen). Library was amplified with KAPA HiFi HotStart Real-time PCR Master Mix (Roche) using indexed primers and sequenced on an Illumina Hiseq 2500 (paired-end 50 bp reads).

### RNA-sequencing with Smart-seq

Smart-seq was performed as described previously^34, 57^. Briefly, total RNA was isolated from naïve and activated B cells with Trizol (ThermoFisher) and the integrity of the RNA was accessed with TapeStation RNA Analysis ScreenTape or Bioanalyzer RNA pico kit (Agilent). 10ng of RNA was reverse transcribed using oligo-dT_30_ VN primer in the presence of Template Switching Oligo (TSO) with SuperScript II reverse transcriptase. cDNA was pre-amplified with IS PCR primers and PCR products were cleaned up with Ampure XP beads. One ng of PCR product was used to generate library using NexteraXT library prep kit (Illumina) and tagmentated DNA was amplified for a 12 cycles PCR and purified with AmpureXP beads. Libraries were sequenced on an Illumina Hiseq 2500 with single-end 50 bp reads.

### Statistical analyses

Statistical analyses and bar plots were performed and plotted with Prism 7 or R (v3.3.3). Bar graph and dot plots shown indicate mean and standard error.

## Materials and Methods --- Bioinformatics analyses

The reference genome used was mm10. Heatmaps and profile plots were generated using DeepTools^58^.

### CMSIP analysis

Paired-end reads (50bp) were mapped to the mouse genome mm10 GRCm38 (Dec. 2011) from UCSC, using BSMAP (V.2.74) (-v 4 -R -n 1 -w 2 -r 0 -q 20 -R -p 8)^59^. Reads that mapped to the spike-in control (Lambda and Puro) were filtered out from the Sam file using awk. Tag directories were created with the remaining reads using makeTagDirectory from HOMER^60^ (-genome mm10 -tbp 1 ‒checkGC). Peaks were called with findPeaks from HOMER (-style histone -o auto -i). Peaks from all samples were merged with mergePeaks from HOMER into a master table. Quantile normalization was applied to all raw counts files and differentially enriched 5hmC regions were identified with edgeR^61^; a *p* adjusted value of ≤ 0.05 was used as a cutoff.

### H3K27Ac ChIP analysis

Single end reads (50bp) were mapped to the mouse genome mm10 GRCm38 (Dec. 2011) from UCSC with Bowtie (V.1.1.2). Reads were sorted and PCR duplicates were removed using SortSam and MarkDuplicates, respectively from Picard Tools (V.2.7.1). Tag directories were created with makeTagDirectory (-genome mm10 ‒checkGC) from HOMER, and peaks were called with findPeaks (-region). Peaks from all samples were merged with mergePeaks from HOMER into a master table. Quantile normalization was applied to all raw counts files and differentially enriched 5hmC regions were identified with edgeR^61^; a *p* adjusted value of ≤ 0.05 was used as a cutoff.

### BATF ChIP analysis

Single end reads (50bp) were mapped to the mouse genome mm10 GRCm38 (Dec. 2011) from UCSC with Bowtie (v1.1.2). Reads were sorted and PCR duplicates were removed using SortSam and MarkDuplicates, respectively from Picard Tools (V.2.7.1). Tag directories were created with makeTagDirectory (-genome mm10 ‒checkGC) from HOMER^60^, and peaks were called with findPeaks (-style factor -o auto).

### Definition of preferentially active enhancers

Preferentially active enhancers (**Supplementary Fig. 1h**) were defined as distal H3K27Ac regions (> 1kb from TSS) that had ATAC-seq and H3K27Ac peaks overlapping in at least 50% of either peak/region; overlapping was calculated with intersectBed -f 0.5 -f 0.5 ‒e (Bedtools v2.26.0). The differential enrichment in an enhancer was called if it contains a differentially enriched H3K27Ac region as well as at least one differentially access region.

### ATAC-seq analysis

Paired-end reads (100 bp) were mapped to the mouse genome mm10 GRCm38 (Dec. 2011) from UCSC using Bowtie 1.0.0 (“-p 8 -m 1 --best --strata -X 2000 -S --fr --chunkmbs 1024.”). Reads that failed this alignment step were filtered for Illumina adapters and low quality using “Trim Galore!” (“--paired --nextera --length 37 --stringency 3 --three_prime_clip_R1 1 --three_prime_clip_R2 1”) and re-mapped using the same parameters. Both mapping results were merged and processed together to remove duplicates using picard-tools-1.94 MarkDuplicates. Mitochondrial and Chromosome Y reads were excluded.

Subnucleosomal fragments were obtained with SAMtools and awk to identify DNA fragments that were less than or equal to 100 nt in length. These fragments were used to call peaks using HOMER (v4.9.1) findPeaks function for each replicate (“-size 500 -region -center -P .1 -LP .1 -poisson .1 -style dnase”) and all the peak sets were merged to generate a global set. Peaks overlapping with ENCODE blacklisted regions^62^ were removed. From each sample, Tn5 insertion sites were obtained by isolation of the initial 9bp of mapped reads^63^ which were used to compute the number of transposase insertions per peak using MEDIPS^64^. Raw reads from all samples were quantile-normalized prior to differential coverage analysis using edgeR without TMM (Trimmed mean of M-values) normalization. Only regions with more than 32 normalized reads across the samples per comparison. Differentially accessible regions were defined by an adjusted *p* value (FDR) lower than 0.05 and a log 2 fold enrichment higher equal than 1.

### OxBS analysis

OxBS-seq reads were mapped to both the mouse genome mm10 GRCm38 (Dec. 2011) from UCSC and the phage Lambda genome (GenBank: J02459.1) using bsmap-2.90 (“-v 15 -w 3 -p 4 -S 1921 -q 20 -A AGATCGGAAGAGC -r 0 -R -V 2”). The mapping results were separated into reads belonging to the mm10 genome and each of the three loci from lambda used for oxidation and conversion efficiency calculation. Methylation calls from lambda-and mm10-derived reads were obtained using bsmap-2.90 function methratio.py (“-u -p -g -i “correct” -x CG,CHG,CHH”). Conversion efficiencies as well as posterior probabilities of methylation, hydroxymethylation and unmodified cytosine were calculated by luxGLM v.0.666 (prior probabilities used for for C, hmC and mC “998,1,1”, “6,2,72” and “1,998,1” respectively)^65^. Following genomic positions from lambda used for oxidation and BS treatment efficiencies: chrLambda:22893-23053 C; chrLambda:23765-23925 hmC; chrLambda:47335-47495 mC.

### WGBS analysis

WGBS reads were mapped to both the mouse genome mm10 GRCm38 (Dec. 2011) from UCSC. Bisulfite conversion efficiency was estimated based on cytosine methylation in non-CpG context. For all the samples the bisulfite conversion efficiency was higher than 0.9996. Duplicated reads caused by PCR amplification were removed by BSeQC (v1.2.0) applying a *p* value cutoff Poisson distribution test in removing duplicate reads (1e-5)^66^. Consequently, a maximum of three stacked reads at the same genomic location were allowed and kept for further analysis. In addition, BSeQC was employed for removing DNA methylation artifacts introduced by end repair during adaptor ligation. Overlapping segments of two mates of a pair were reduced to only one copy to avoid considering the same region twice during the subsequent DNA methylation quantification. To estimate CpG DNA methylation at both DNA strands, methratio.py script was executed from BSMAP (v2.90) (-u -r -z -g -i “correct” -x CG). To identify differentially methylated cytosines and regions (DMCs and DMRs), a naïve B cells dataset and was compared to a activated B cell replicate using RADmeth methpipe-3.4.2 (adjust -bins 1:100:1; merge -p 0.05)^67^.

### RNA-seq analysis

RNA-seq samples at four different time-points collected from WT and DKO conditions were first mapped to the mouse genome mm10/GRCm38 using both Hisat2^68^ (“--no-mixed --no-discordant --add-chrname ‒dta”) and Tophat2^69^ (“--no-novel-juncs”) alignment programs separately. Aligned bam files obtained from both the programs were further used to generate the Hisat2- and Tophat2-specific counts using HTseq-count program^70^ (default parameters). Hisat2- and Tophat2-specific count files at each time point for WT and DKO conditions were then used to identify the differentially expressed genes (FDR < 0.05) between matching time points using edgeR program^61^. Potential batch effects were removed using svaseq program^71^. Finally, the common differentially expressed genes obtained from both Hisat2- and Tophat2-specific list were used to perform the downstream analysis.

### Genome-browser track generation for ChIP-seq

ChIP-seq results from Tet2, Ig control, E2A, PU.1, p300, and GCN5 were processed as follow to generate the genome browser tracks. Fastq files were mapped to mm10 reference genome with Bowtie 2 (v2.1.0) with “-very-sensitive”. The mapped SAM files were converted to BAM using Samtools (v1.7) view ‒h ‒F 4, and duplicates were removed using Picard (v2.7.1). BigWig files were made by first generating a BedGraph files from the filtered Bam files using Bedtools (v2.26.0) genomecov followed by bedGraphToBigWig (v4) with read counts normalized to 10,000,000 reads.

### Miscellaneous analyses of regions

For distance between regions to the closest TSS was analyzed with HOMER software with “annotatePeaks.pl ‒ annStats”. Overlap between regions was analyzed by “bedtools intersect” with no requirement for the degree of overlapping. The degree of significance for overlap between superenhancers and test regions was estimated by Fisher exact test using “bedtools fisher”.

### Time-series analysis

For a unified analysis of the RNA-seq time-course data (0hr to 96hr) from WT samples, TCseq package^72^ was used on the combined RNA-seq read counts, obtained after applying Tophat2^69^ and Hisat2^68^ alignment programs (Described in the previous RNA-seq analysis part). TCseq utilizes GLM method implemented in edgeR package^61^ to detect the differential events in gene expression. Differential analysis was performed between “0hr” to the rest of the time points, and the significant differential events were extracted whenever a log2FC > 2 or < −2 and FDR < 0.05 criteria was satisfied. To detect the temporal pattern of the differential gene expressions (RPKM values), a soft clustering algorithm implemented in TCseq program was applied (“algo = ‘cm’, k = 6, standardize = TRUE”). Finally, the differential genes were assigned to a cluster (C1-C6) representing a specific temporal pattern of expression, if the membership probability of the genes to a cluster is 0.5 or more.

### Published datasets

Naïve H3K4me1 (0h): SRR1535686, SRR1535685. Activated H3K4me1 (48h): SRR1014530. SRR1087900. Naive WGBS (0h): SRR1003257. Activated WGBS (48h): SRR1020523. Naive PU1 (0h): SRR2976278. Activated PU1 (48h): SRR1014532. Naïve DSG control (0h): SRR3158132. Activated E2A DSG anti-CD40/IL-4 (48h): SRR3158146. Naïve Brg1 (0h): SRR3619348. Naïve Chd4 (0h): SRR3619349. Naïve Gcn5 (0h): SRR3619350. Naïve p300 (0h): SRR3619356. Activated Brg1 (24h): SRR3619334. Activated Chd4 (24h): SRR3619335. Activated Gcn5 (24h): SRR3619336. Activated p300 (24h): SRR3619342.

